# Chemogenetic modulation of luciferase emission color for imaging and sensing

**DOI:** 10.1101/2025.11.21.689291

**Authors:** Horgan Manirakiza, Yuriy Shpinov, Amandine Gontier, Lina El Hajji, Ludovic Jullien, Arnaud Gautier

## Abstract

Bioluminescent luciferases have emerged as powerful tools for bioimaging, enabling to image biological systems without external excitation light, reducing thus phototoxicity and eliminating background autofluorescence. Advanced imaging requires luciferases that deliver high photon output for enhanced sensitivity, tunable emission colors for multicolor imaging, and red-shifted emission for effective deep tissue imaging. Here, we introduce LumiFAST, a small tunable luciferase engineered by fusing the bright blue-light emitting NanoLuc with the tunable chemogenetic fluorescent reporter pFAST. pFAST binds and stabilizes the fluorescent state of a variety of synthetic fluorogenic chromophores (also called fluorogens). Its proximity with NanoLuc leads to efficient bioluminescence resonance energy transfer (BRET), enabling customizable red-shifted emission. Thanks to the small size of pFAST, LumiFAST maintains a compact structure, while its modular design allows emission color to be tuned from cyan to green, yellow, orange and red simply by changing the fluorogen. Systematic optimization of the fusion topology and linker length yielded an optimal variant with apparent BRET efficiencies reaching up to 90 %. The red-shifted emission of LumiFAST enables dual-color microscopy imaging when used alongside NanoLuc and allows imaging through thick scattering media. Beyond imaging, our insights into the structural factors governing efficient BRET allowed us to engineer biosensors based on NanoLuc and pFAST for the visualization of protease activity and protein-protein interactions in live cells.

## INTRODUCTION

Optical reporters have become indispensable tools for investigating cellular processes. They can be targeted to specific cells to allow the tracking of individual cells in a population or used to visualize subcellular structures or specific cellular proteins. Although fluorescent proteins are widely used, bioluminescent luciferases are attracting increasing attention in the bioimaging field^1,2^. Unlike fluorescent proteins, luciferases do not require external excitation light – they produce light through catalytic oxidation of luciferin substrates – limiting thus phototoxicity and reducing background autofluorescence.

The engineering of luciferases has primarily focused on two objectives: (i) increasing photon output to enhance spatiotemporal resolution and allow luciferase to compete with fluorescence techniques, and (ii) shifting emission to longer wavelengths (red-shifting) to enable deeper tissue penetration and reduce light scattering, thereby improving in vivo imaging. However, achieving high photon output and red-shifted emission simultaneously has proven challenging. For example, Firefly luciferase, the first bioluminescent reporter, emits yellow light at 560 nm, allowing intravital imaging in small animals^3^. However, its relatively dim bioluminescence limits its application in microscopy. Conversely, NanoLuc, an engineered luciferase of 19 kDa derived from the shrimp *Ophlophorus*^4^, displays a 100-fold higher photon output when used with the luciferin furimazine^5^, making it suitable for microscopy imaging. However, NanoLuc emits blue light peaking at 460 nm, restricting its effectiveness for in vivo imaging due to limited penetration of light into tissues.

The emission of NanoLuc has been recently shifted to higher wavelengths by fusing NanoLuc to fluorescent proteins to induce bioluminescence resonance energy transfer (BRET). This approach led to the development of the enhanced NanoLantern series, created by fusing NanoLuc to fluorescent proteins spanning a broad spectrum from cyan to red, enabling efficient five-color imaging in live cells^6^. Building on these advances, the enhanced NanoLantern series was further diversified to include 20 distinct colors by adding a second fluorescent protein, producing additional hues through dual-acceptor BRET^7^. In parallel, the Antares variant was engineered by fusing NanoLuc to CyOFP1, a large Stokes-shift red fluorescent protein, enabling efficient in vivo imaging^8^. Furthermore, luciferases with tunable emission colors have been generated by fusing NanoLuc to HaloTag, a self-labeling protein that can be labeled with various synthetic fluorophores, thus enabling precise modulation of the bioluminescence emission color^9^.

While these red-shifted bioluminescent constructs based on fluorescent proteins or HaloTag have proven effective, their significant larger size compared to NanoLuc alone compromise one of NanoLuc’s main advantages: its compactness as reporter. In this work, we introduce the tunable luciferase LumiFAST, an optimized fusion of NanoLuc and the compact 14 kDa chemogenetic reporter pFAST, which binds and stabilizes the fluorescent states of a range of fluorogenic chromophores (also called fluorogens)^10^ (Examples are shown in **Supplementary Figure 1** and **Supplementary Table 1**). The interest of using pFAST lies in its small size and its modularity: its small size allows LumiFAST to remain relatively small (33 kDa), and its modularity enables modulation of the bioluminescence emission wavelength from cyan to green, yellow, orange or red by selecting different fluorogens (**Figure 1a,b**). Previous work has shown that directly fusing pFAST to NanoLuc enables to modulate bioluminescence emission through BRET^11^. Here, through systematic optimization of the fusion topology and linker length, we have developed an enhanced variant, LumiFAST, which achieves apparent BRET efficiencies up to 90 %. The red-shifted emission of LumiFAST enables dual-color microscopy imaging when combined with standard NanoLuc and allows imaging through deep tissues ex vivo. Beyond imaging applications, our understanding of the structural requirements for efficient BRET also enabled us to leverage NanoLuc and pFAST for creating intracellular biosensors to visualize protease activity and protein-protein interactions in live cells.

**Figure 1.**
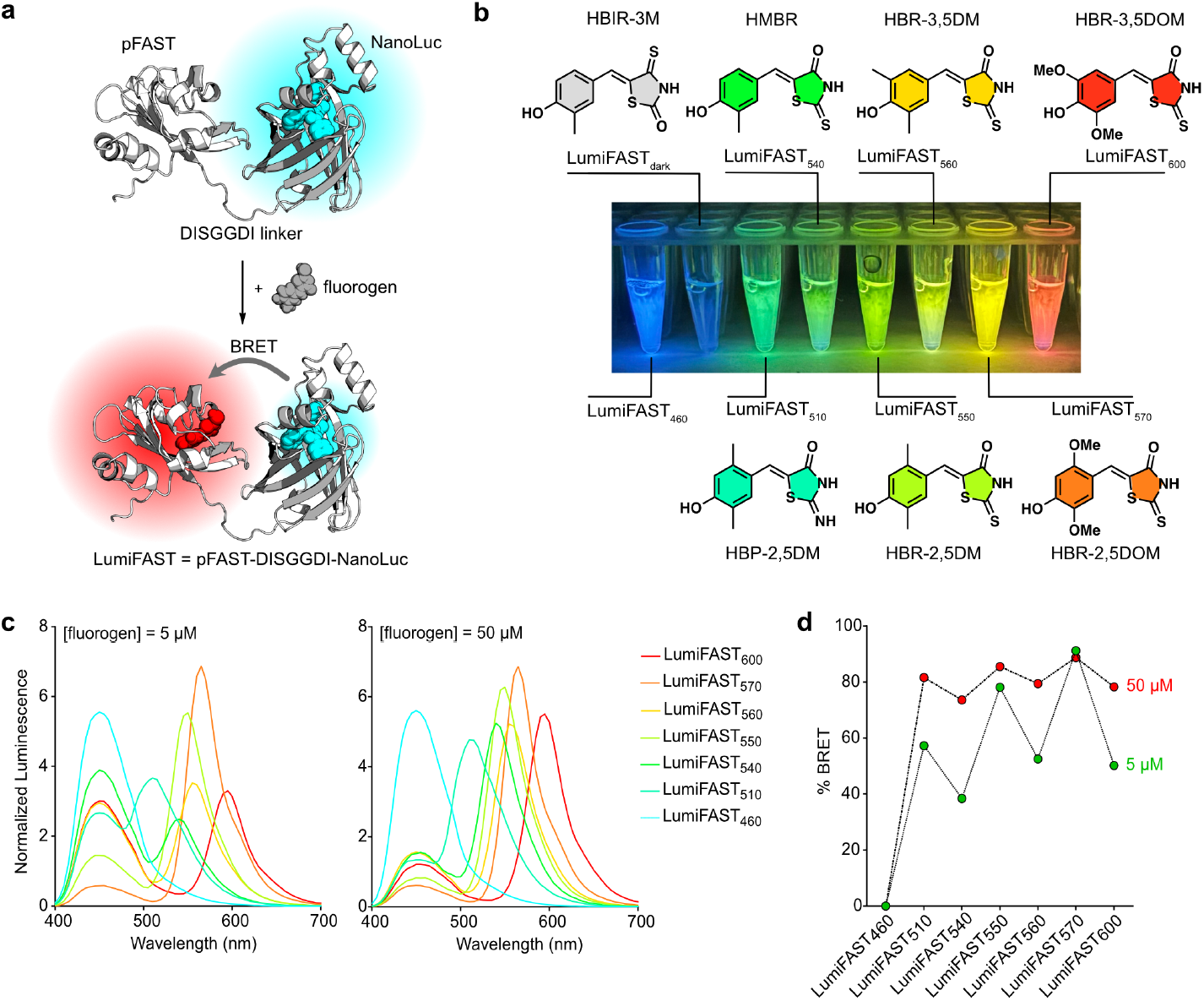
LumiFAST, a tunable chemogenetic luciferase. **a**. Principle of LumiFAST. The model of LumiFAST was generated with Alphafold3, positioning the fluorogen and furimazine in Pymol. **b**. The choice of fluorogen enables to tune the emission color of LumiFAST. Smartphone photographs of solutions of LumiFAST (50 nM) with various fluorogens (5 μM) and with final 250-fold dilution of NanoGlo Furimazine substrate (Promega). **c**. Bioluminescence emission spectra of LumiFAST (50 nM) with various fluorogens (5 μM or 50 μM) and with final 250-fold dilution of NanoGlo Furimazine substrate (Promega). The spectra are the average of three independent measurements. Spectra were normalized by the total bioluminescence intensity (area normalization). **d**. Apparent BRET efficiency of the different chemogenetic luciferases shown in panel **c** in function of the fluorogen concentration. The data represents the mean of three independent measurements.

## RESULTS

### Design and optimization of LumiFAST

Given that BRET efficiency is highly dependent on both the orientation and distance between proteins, we initially compared N- and C-terminal fusions of NanoLuc (as well as its circular permutation ^CP^NanoLuc obtained by fusing the original N- and C-termini of NanoLuc with a (GGTGGS)_2_ linker and introducing new N- and C-termini in position 66 and 65^9^) with pFAST (and its circular permutation ^CP^pFAST obtained by fusing the original N- and C-termini of pFAST with a GGSGGSGGSGG linker and introducing new N- and C-termini in position 115 and 114 (see **Supplementary Figure 2** for characterization)). All constructs used a 5-amino-acid DISGG flexible linker. This initial screen showed that the fusion pFAST-DISGG-NanoLuc provided the highest apparent BRET efficiency when using the fluorogen HBR-3,5DOM (λ_abs_ 520 nm, λ_em_ 600 nm) (**Supplementary Figure 3a**). To further evaluate how the linker length impacts BRET efficiency, we generated nine additional fusions, each containing a linker of 0, 3, 5, 6, 7, 8, 9, 10 or 11 amino acids. When tested with various fluorogens – HBR-2,5DM (λ_abs_ 495 nm, λ_em_ 550 nm), HBR-2,5DOM (λ_abs_ 510 nm, λ_em_ 570 nm) and HBR-3,5DOM (λ_abs_ 520 nm, λ_em_ 600 nm) – the construct with 7-amino-acid DISGGDI linker consistently yielded the highest apparent BRET efficiency, reaching 80 %, 90 % and 50 %, respectively, when using 5 μM of fluorogen (**Supplementary Figure 3b**,**c**). Alongside these optimizations, we also explored the insertion of pFAST or ^CP^pFAST into NanoLuc or ^CP^NanoLuc and vice versa (**Supplementary Figure 1d**). However, none of these alternative configurations surpassed the apparent BRET efficiency of the optimized pFAST-DISGGDI-NanoLuc, which we, hereafter, refer to as LumiFAST.

We next examined the spectral properties of LumiFAST using a range of fluorogens including HBP-2,5DM (λ_abs_ 460 nm, λ_em_ 510 nm), HMBR (λ_abs_ 480 nm, λ_em_ 540 nm), HBR-2,5DM (λ_abs_ 495 nm, λ_em_ 550 nm), HBR-3,5DM (λ_abs_ 500 nm, λ_em_ 560 nm), HBR-2,5DOM (λ_abs_ 510 nm, λ_em_ 570 nm), HBR-3,5DOM (λ_abs_ 520 nm, λ_em_ 600 nm). These fluorogens were specifically selected for their diverse emission wavelengths and their strong absorption in the blue region when bound to pFAST, which is optimal for efficient BRET with NanoLuc. By pairing LumiFAST with each of these fluorogens (at 5 μM), we successfully generated luciferase variants exhibiting distinct emission peaks at 510, 540, 550, 560, 570 and 600 nm (**Figure 1b,c**), and apparent BRET efficiency of 57 %, 38 %, 78 %, 53 %, 92% and 51% (**Figure 1d**). Interestingly, when using the dark fluorogen HBIR-3M (λ_abs_ 495 nm), which forms a non-fluorescent assembly with pFAST, we also observed a marked quenching of bioluminescence (**Supplementary Figure 4**), confirming effective energy transfer to this “dark” acceptor. For clarity, we designate each LumiFAST:fluorogen variant by its emission wavelength as a subscript: for example, LumiFAST paired with HBR-2,5DOM (emitting at 570 nm) is referred to as LumiFAST_570_.

It is important to highlight that the BRET efficiency values reported here are considered “apparent” as they depend on the fluorogen concentration used. In our experiments, employing 5 μM of fluorogen maximizes the formation of the pFAST:fluorogen complex (**Supplementary Figure 1** and **Supplementary Table 1**), but at this concentration, a fraction of NanoLuc’s blue emission is absorbed by the free, unbound fluorogen via an inner filter effect (**Supplementary Figure 5**). This absorption reduces the observed NanoLuc emission peak and artificially increases the calculated BRET efficiency, resulting in an overestimation of the true BRET efficiency. When the fluorogen concentration is increased tenfold to 50 μM, the inner filter effect becomes even more pronounced (**Supplementary Figure 5**), further increasing the apparent BRET efficiency (**Figure 1c,d**). However, this also leads to a substantial reduction in the total bioluminescence intensity. A comprehensive analysis of how both BRET efficiency and bioluminescence intensity change with fluorogen concentration can be found in **Supplementary Figure 6**. Based on these findings, we determined that using 5 or 10 μM of fluorogen offers the best compromise between achieving high apparent BRET efficiency and maintaining high photon output for imaging applications.

During the course of this work, Prescher et al. introduced BREAKFAST (Bioluminescence Resonance Energy mAKe over with a Fluorescence-Activating absorption-Shifting Tag), a luciferase with tunable emission generated by directly fusing pFAST and NanoLuc (pFAST-NanoLuc)^11^ – a design identical to our construct without linker. To provide a comprehensive comparison, we evaluated the performances of LumiFAST and BREAKFAST side by side using various fluorogens at 5 and 50 μM (**Supplementary Figures 7** and **8**). Consistent with our findings on the impact of the linker length on the BRET efficiency, LumiFAST consistently outperformed BREAKFAST. Specifically, at 5 μM of fluorogen, LumiFAST exhibited higher apparent BRET efficiency than BREAKFAST with HBP-2,5DM, HMBR, HBR-2,5DM, HBR-3,5DM, HBR-2,5DOM and HBR-3,5DOM by 1.9-, 2.4-, 1.8-, 1.5-, 1.8- and 3.8-fold respectively (**Supplementary Figure 8**). This trend was maintained at 50 μM of fluorogen, conditions previously used by Prescher et al. for characterizing BREAKFAST^11^, where LumiFAST showed superior apparent BRET efficiencies compared to BREAKFAST with the same set of fluorogens (1.9-, 1.9-, 1.8-, 1.5-, 1.9- and 3-fold higher respectively) (**Supplementary Figure 8**).

### Imaging of LumiFAST in cultured mammalian cells

We next showed that LumiFAST could be an efficient bioluminescent reporter in cultured mammalian cells. For bioluminescence imaging, we used a fluorescence microscope equipped with an EMCCD camera, switching off the excitation laser to exclusively detect bioluminescence signal, and using a 10 s exposure time to ensure optimal sensitivity. Under these conditions, HeLa cells expressing LumiFAST and treated with HBR-2,5DM (to form LumiFAST_550_), HBR-2,5DOM (to form LumiFAST_570_) and HBR-3,5DOM (to form LumiFAST_600_) in the presence of furimazine displayed homogenous bioluminescent signals throughout the cells (**Figure 2**). This bioluminescence distribution closely mirrored the fluorescence patterns observed when imaging LumiFAST fluorescence in the absence of furimazine. To further characterize the spectral properties of the different LumiFAST variants, we applied a series of emission filters. In the absence of fluorogen, LumiFAST showed maximal emission in the blue channel (420-480 nm), while LumiFAST_550_ displayed maximal emission in the green channel (500-550 nm), LumiFAST_570_ in the red channel (580-625 nm) and LumiFAST_600_ in the red (580-625 nm) and far-red channel (670-780 nm). These results were consistent with the specific maximal emission wavelengths of each LumiFAST:fluorogen combination.

**Figure 2.**
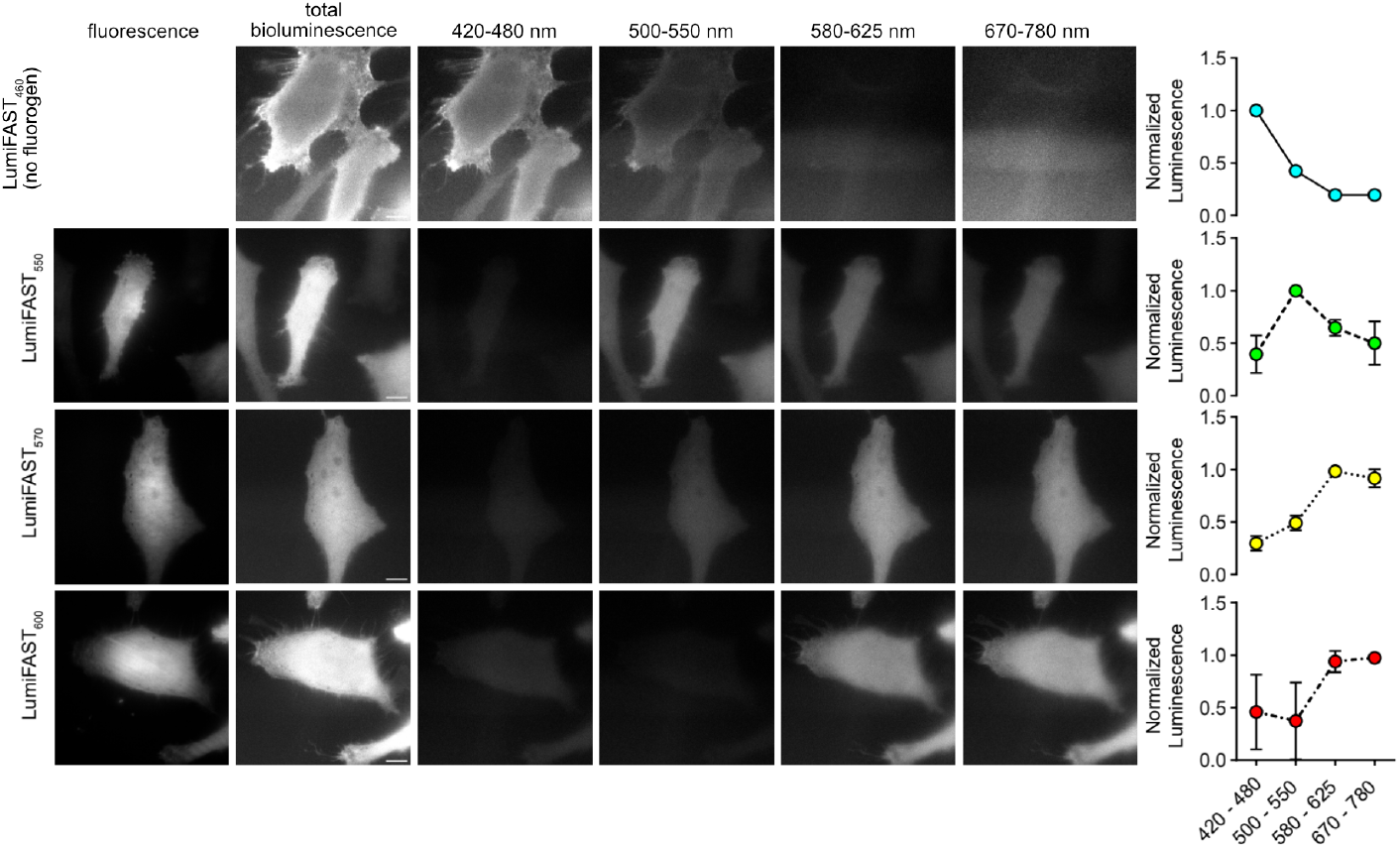
LumiFAST for tunable bioluminescence imaging in live mammalian cells. Live HeLa cells expressing LumiFAST were treated with no fluorogen or with 10 μM of HBR-2,5DM (to form LumiFAST_550_) or HBR-2,5DOM (to form LumiFAST_570_) or HBR-3,5DOM (to form LumiFAST_600_). Cells were first imaged by fluorescence microscopy (images on the left). Next, cells were additionally treated with final 50-fold dilution of NanoGlo Furimazine substrate (Promega), and imaged by bioluminescence microscopy using different spectral channels to characterize the red shift in emission (exposure time 10 s). Representative micrographs from three independent experiments. Scale bars 10 μm. Graphs on the right show relative bioluminescence intensities (mean ± SD of three independent experiments) in the different spectral channels.

We further demonstrated the versatility of LumiFAST for imaging proteins in various cellular localizations. By creating fusions of LumiFAST to the nuclear histone H2B, to Lyn11 membrane targeting sequence and to the outer mitochondrial protein Tom20, we confirmed that each fusion accurately localized to its intended cellular localization in live HeLa cells, showing that LumiFAST does not perturb the normal localization of these proteins (**Figure 3**). Consistent with our earlier observations, the spatial pattern of bioluminescence closely matched that of the corresponding fluorescence signals. To enhance image quality, we applied the Noise2Void 2D deep-learning method, which denoises 2D microscopy images. We trained the neural network using high-quality fluorescence micrographs of cells expressing LumiFAST, then applied the trained network to our bioluminescence micrographs, resulting in a significantly improved signal-to-noise ratio (**Supplementary Figure 9**).

**Figure 3.**
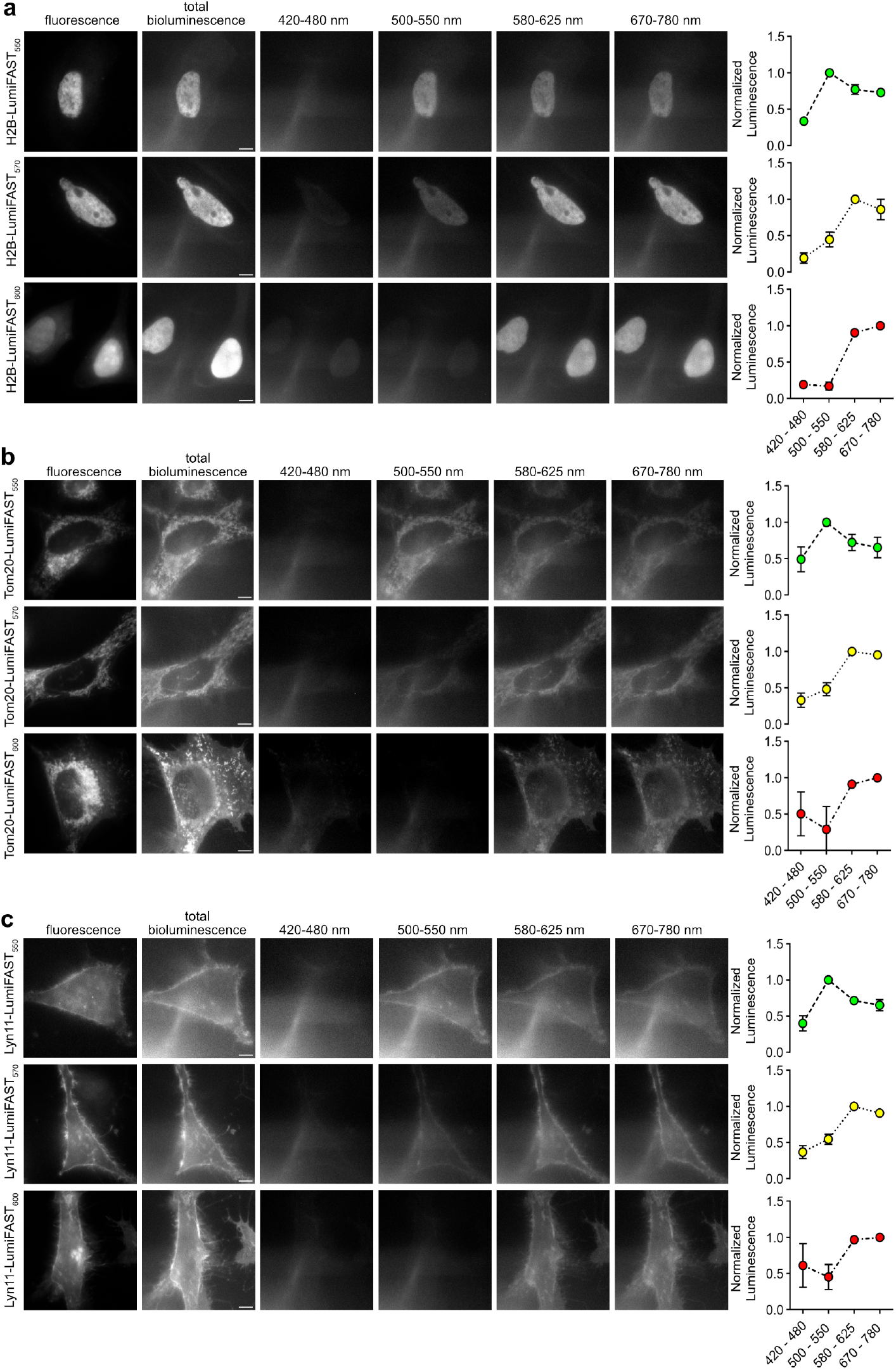
Selective bioluminescence imaging of LumiFAST in various cellular localizations in live mammalian cells. Live HeLa cells expressing H2B-LumiFAST (**a**), Tom20-LumiFAST (**b**), Lyn11-LumiFAST (**c**) were treated with 10 μM of HBR-2,5DM (to form LumiFAST_550_) or HBR-2,5DOM (to form LumiFAST_570_) or HBR-3,5DOM (to form LumiFAST_600_). Cells were first imaged by fluorescence microscopy (images on the left). Next, cells were additionally treated with final 50-fold dilution of NanoGlo Furimazine substrate (Promega), and imaged by bioluminescence microscopy using different spectral channels to characterize the red shift in emission (exposure time 10 s). Representative micrographs from three independent experiments. Scale bars 10 μm. Graphs on the right show relative bioluminescence intensities (mean ± SD of three independent experiments) in the different spectral channels.

Next, we demonstrated that LumiFAST could be effectively combined with NanoLuc for two-color bioluminescence imaging. We performed bioluminescence imaging on HeLa cells co-expressing LumiFAST and NanoLuc, each targeted to distinct subcellular localizations (**Figure 4**). Following treatment with 10 μM HBR-2,5DOM (to form LumiFAST_570_) and furimazine, we observed distinct blue bioluminescence from regions expressing NanoLuc, and red bioluminescence from areas expressing LumiFAST_570_. These results clearly showed the ability to perform simultaneous two-color bioluminescence imaging using NanoLuc and LumiFAST.

**Figure 4.**
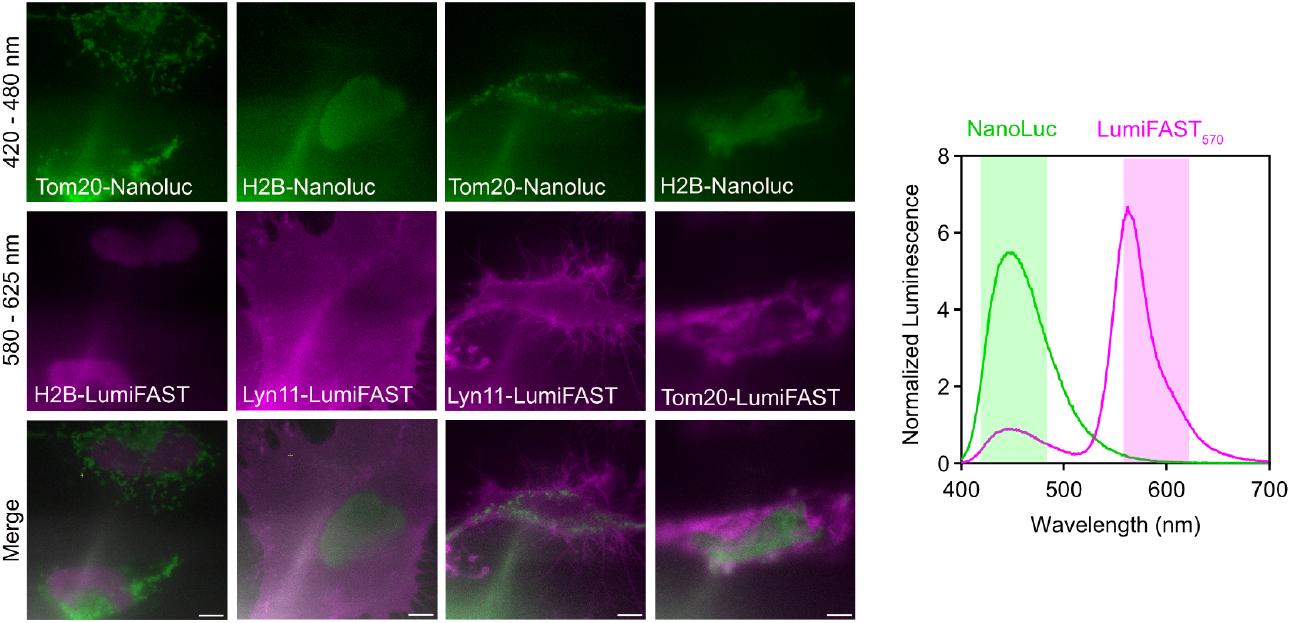
Selective two-color bioluminescence imaging of LumiFAST and NanoLuc in live mammalian cells. Live HeLa cells expressing LumiFAST and NanoLuc targeted to different subcellular localization were treated with 10 μM of HBR-2,5DOM (to form LumiFAST_570_) and with final 50-fold dilution of NanoGlo Furimazine substrate (Promega). Cells were imaged by bioluminescence microscopy using 420-480 nm filter (to detect NanoLuc bioluminescence) and 580-625 nm filter (to detect LumiFAST_570_ bioluminescence) (exposure time 10 s). Representative micrographs from three independent experiments. Scale bars 10μm.

### Imaging of LumiFAST through thick, highly scattering samples

We next assessed the suitability of the LumiFAST variants for deep-tissue imaging. To do so, we first imaged solutions of the LumiFAST variants (25 nM) through a 1 cm agar-based tissue phantom designed to mimic the light absorption and scattering properties of soft tissues. In this experimental set-up, LumiFAST_570_ and LumiFAST_600_ emerged as the most effective for imaging through thick, highly scattering samples (**Figure 5a**), which aligns with their red-shifted emission profiles. To further validate these findings, we imaged LumiFAST solutions (25 nM) through 1 cm thick chicken breast tissue. Consistent with the phantom experiments, LumiFAST_570_ and LumiFAST_600_ again demonstrated superior performance (**Figure 5b**), underscoring their strong potential for deep-tissue imaging applications.

**Figure 5.**
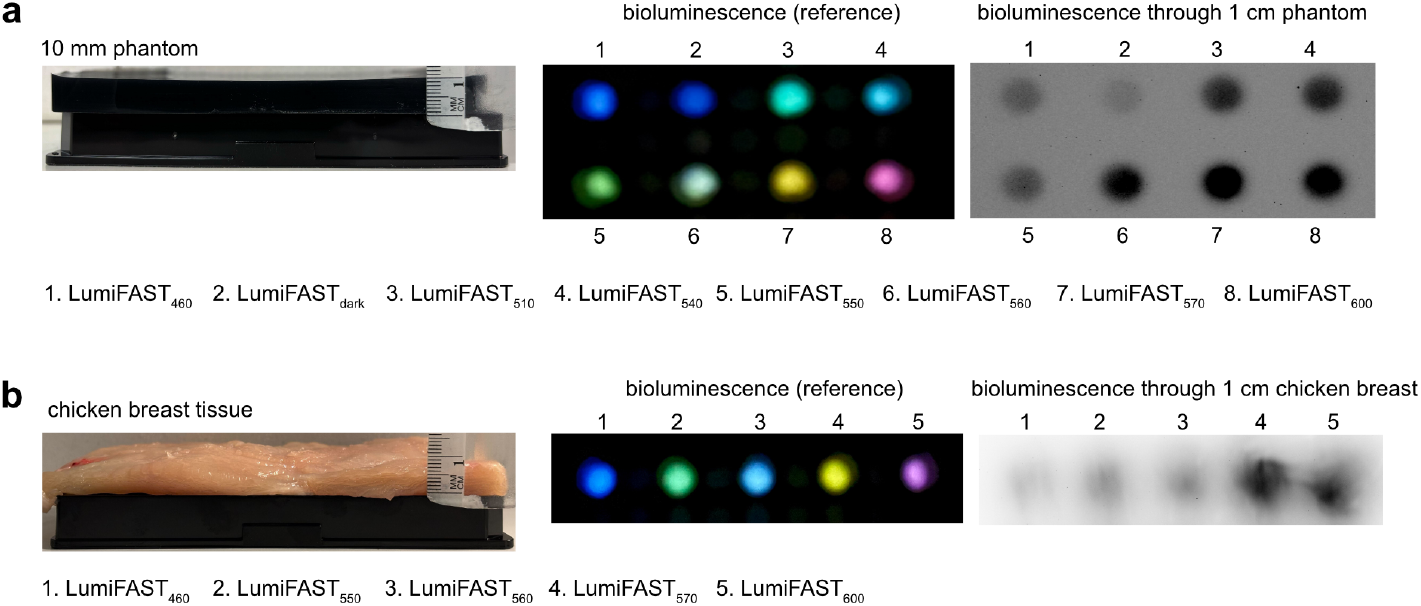
Imaging of LumiFAST through thick scattering samples. **a** Solutions of LumiFAST protein (25 nM), fluorogens (5 μM) with final 250-fold dilution of NanoGlo Furimazine substrate (Promega) in a multiwell plate were imaged through a 1 cm tissue phantom. The color image shows the color of the different LumiFAST variants in absence of the phantom (photographs taken in the dark with a smartphone). The black and white image shows the bioluminescence of the different spectral LumiFAST variants through the 1 cm tissue phantom (photographs taken with a Gel Imager). **b** Solutions of LumiFAST protein (25 nM), fluorogens (5 μM) with final 250-fold dilution of NanoGlo Furimazine substrate (Promega) in a multiwell plate were imaged through a 1 cm chicken breast tissue. The color image shows the color of the different LumiFAST variants in absence of the phantom (photographs taken in the dark with a smartphone). The black and white image shows the bioluminescence of the different spectral LumiFAST variants through the 1 cm chicken breast tissue (photographs taken with a Gel Imager).

### Design of BRET-based protease sensors

The high BRET efficiency observed in LumiFAST led us to explore the development of protease sensors by inserting a protease cleavage site as linker between pFAST and NanoLuc. In theory, proteolytic cleavage at this site would physically separate the two proteins, resulting in a measurable decrease in BRET signal. As a proof-of-concept, we engineered a sensor for the Tobacco Etch Virus protease (TEVp), a widely used tool in biological and biotechnological applications^12–15^. To construct this TEVp sensor, we replaced the original linker of LumiFAST by the sequence SENLYFQS, which is specifically recognized and cleaved by TEVp^12^. This design preserved a high BRET efficiency in vitro (**Figure 6a-c** and **Supplementary Figure 10**). However, when we incubated the purified recombinant sensor with TEVp overnight, we saw no change in BRET efficiency (**Figure 6c** and **Supplementary Figure 9**). Analysis by SDS-PAGE further revealed that the sensor remains uncleaved under these conditions (**Figure 6b**), consistent with the absence of any BRET change. We hypothesized that the 8-amino-acid long sequence used as linker was too short to allow TEVp sufficient access to its recognition sequence within the fusion protein. To address this, we extended the linker length to 12 and 19 amino acids by adding additional residues flanking the SENLYFQS site. While increasing the linker length resulted in a progressive decrease in BRET efficiency – the longer the linker, the greater the reduction – both extended systems exhibited a complete loss of BRET upon incubation with the TEVp, indicating successful proteolytic cleavage (**Figure 6c** and **Supplementary Figure 10**). Analysis by SDS-PAGE confirmed that the two sensor variants were efficiently cleaved by TEVp in vitro (**Figure 6b**), validating our design strategy for creating BRET-based protease sensors.

We further characterized the system with a 12-amino-acid linker in live mammalian cells (**Figure 6d**). In HeLa cells co-expressing both the sensor and TEVp and treated with HBR-2,5DOM, we detected only blue bioluminescence in agreement with complete cleavage by TEVp. In contrast, HeLa cells expressing the sensor alone (without TEVp) and treated with HBR-2,5DOM displayed high red bioluminescence, indicating the system remained intact. These findings demonstrate that our strategy enables robust and specific detection of protease activity in living cells. Importantly, the design is generalizable and could be adapted for monitoring a wide range of proteases.

**Figure 6.**
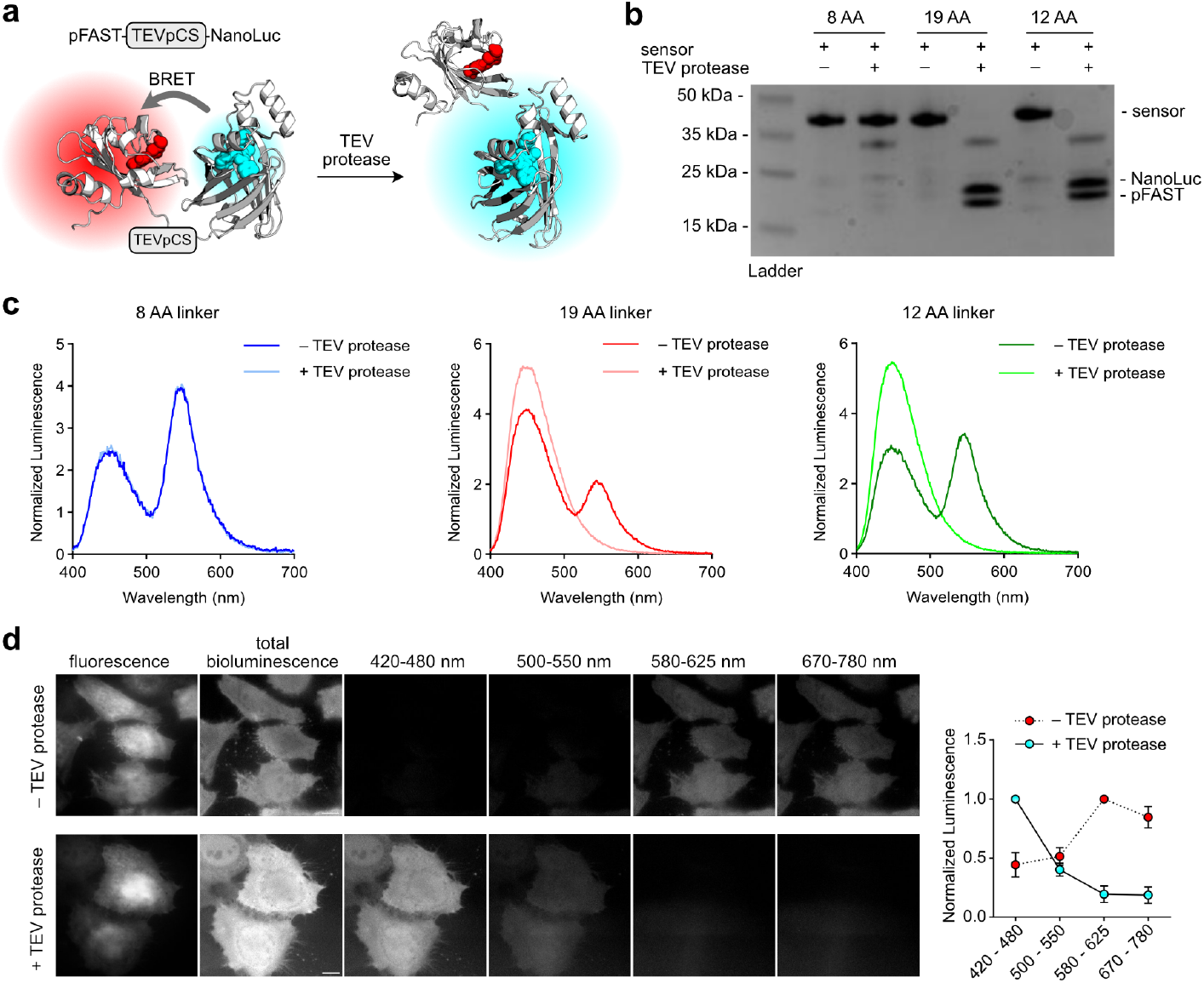
BRET-based sensing of protease activity. **a** Principle of BRET-based TEV protease sensor. **b** SDS-PAGE of sensor solution (20 μM) incubated or not with TEV protease overnight. Different linker lengths were tested: 8, 12 and 19 amino acid long linkers. **c** Bioluminescence emission spectra of the three sensor candidates (50 nM) (HBR-2,5DOM 5 μM with 250-fold dilution of NanoGlo Furimazine substrate (Promega)) before and after treatment with the TEV protease. Spectra were normalized by the total bioluminescence intensity (area normalization). **d** Live HeLa cells expressing the 12 AA linker TEV sensor (Top row) or co-expressing the 12 AA linker TEV sensor and the TEV protease (bottom row) were treated with HBR-2,5DOM (10 μM) and imaged first by fluorescence microscopy. Then, cells were treated with final 50-fold dilution of NanoGlo Furimazine substrate (Promega) and imaged by bioluminescence microscopy in different spectral channels, enabling to monitor the activity of the TEV protease. Representative micrographs of three independent experiments. Scale bars 10 μm. The graph on the right shows the bioluminescence intensity (mean ± SD of three independent experiments) in the different spectral channels.

### Detection of protein-protein interactions

Encouraged by these results, we further investigated the use of pFAST and NanoLuc for the detection of protein-protein interactions. This application extends beyond the design of LumiFAST, leveraging our comprehensive understanding of the system’s modularity. pFAST and NanoLuc can be fused to two proteins, allowing to probe their proximity by monitoring BRET efficiency. As model interaction, we used FKBP (FK506-binding protein) and FRB (FKBP-rapamycin binder), two proteins known to interact specifically in presence of the small molecule rapamycin^16^. To systematically evaluate all possible configurations, we constructed eight fusions by tethering pFAST and NanoLuc to either the N- or C-terminus of FRB and FKBP, respectively, using a DISGGDIS linker. In vitro analysis revealed that the pairing of pFAST-DISGGDIS-FRB and FKBP-DISGGDIS-NanoLuc produced the highest BRET efficiency upon rapamycin addition (**Supplementary Figure 11a**). We then optimized the system by varying the linker length (0, 2 or 8 amino acids) between the fusion partners. The highest BRET efficiency in the presence of rapamycin was achieved when no linker was used between pFAST and FRB and between FKBP and NanoLuc (**Supplementary Figure 11b** and **Figure 7a,b**). Under these optimized conditions, the highest apparent BRET efficiency in the presence of rapamycin approached the maximal values obtained with LumiFAST, indicating a highly favorable molecular arrangement. Furthermore, in vitro bioluminescence titration with varying concentrations of rapamycin allowed to determine an effective rapamycin concentration for half maximal BRET of about 10 nM (**Figure 7c**), closely matching previously reported value^16^. These promising in vitro results were validated in live cells (**Figure 7d**). HeLa cells co-expressing pFAST-FRB and FKBP-NanoLuc, displayed only blue bioluminescence in the absence of rapamycin, in agreement with the absence of interaction of FRB and FKBP. Upon rapamycin addition, we observed a pronounced red-shift in bioluminescence, indicating a successful chemically induced interaction of FRB and FKBP. Altogether, these experiments demonstrate that NanoLuc and pFAST can be effectively combined to monitor protein-protein interactions in live cells with high specificity and sensitivity, broadening the potential applications of this bioluminescent BRET-based system.

**Figure 7.**
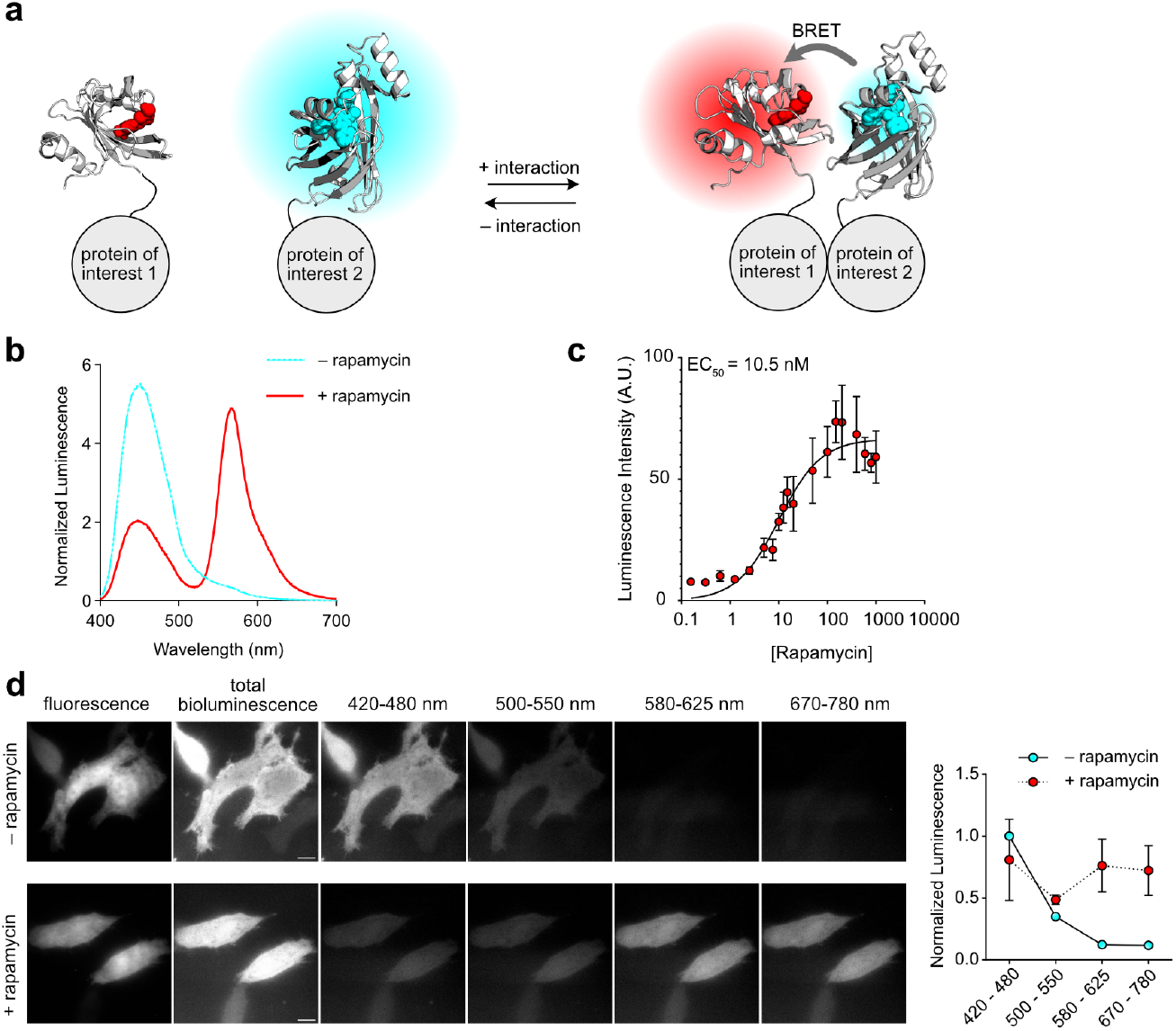
BRET-based sensing of protein-protein interactions. **a** Principle. **b** Bioluminescence emission spectra of solutions of pFAST-FRB (50 nM), FKBP-NanoLuc (25 nM), HBR-2,5DOM (5 μM) with and without rapamycin (100 nM) and with final 250-fold dilution of NanoGlo Furimazine substrate (Promega). Spectra were normalized by the total bioluminescence intensity (area normalization). **c** In vitro evolution of the bioluminescence intensity in function of the rapamycin concentration. **d** Live HeLa cells expressing pFAST-FRB and FKBP-NanoLuc labeled with HBR-2,5DOM (10 μM) were treated with or without rapamycin. Cells were first imaged by fluorescence microscopy (micrographs on the left). Then, cells were treated with final 50-fold dilution of NanoGlo Furimazine substrate (Promega) and imaged by bioluminescence microscopy in different spectral channels. Representative micrographs of three independent experiments. Scale bars 10 μm. The graph on the right shows the bioluminescence intensity (mean ± SD of three independent experiments) in the different spectral channels.

## DISCUSSION

In this study, we report the development of LumiFAST, a compact and highly versatile luciferase platform with tunable emission color, which enables the imaging of fusion proteins in various systems. LumiFAST was engineered by fusing pFAST – a versatile chemogenetic fluorescent reporter whose absorption and emission properties can be precisely tuned by using different fluorogenic chromophores – and NanoLuc, a blue-light-emitting luciferase known for its exceptional photon output. The emission color of LumiFAST can be adjusted through efficient bioluminescent resonance energy transfer (BRET) between NanoLuc and the pFAST:fluorogen complex. As pFAST is a small protein (just 14 kDa), the entire LumiFAST construct is about 33 kDa, making it the smallest BRET-based red-shifted luciferase reported to date.

This modular design allows for straightforward color tuning of LumiFAST – from cyan to green, yellow, orange and red – by simply selecting different fluorogens for pFAST, thus enabling a wide range of applications. Through systematic optimization of both the fusion topology and the linker length connecting the BRET partners, we achieved maximal BRET efficiency and optimal spectral tuning. Compared to BREAKFAST, a previously described pFAST-NanoLuc fusion by Prescher et al.^11^, LumiFAST displays a substantial increase (about 2 to 4-fold at 5 μM fluorogen) in BRET efficiency. This improvement results from a comprehensive investigation of linker length, underscoring the importance of precise engineering in enhancing BRET-based imaging system.

In this study, we show that LumiFAST is an efficient molecular tool for imaging fusion proteins in diverse cellular localizations using bioluminescence microscopy. Its red-shifted emission and optimized BRET efficiency enables to combine it with NanoLuc for dual color imaging through simple spectral discrimination. Looking ahead, LumiFAST and NanoLuc could be further combined with enhanced NanoLantern variants^6^ or with Antares^8^ for highly multiplexed bioluminescence imaging. Furthermore, the use of red-shifted fluorogens enables to generate luciferases that strongly emit in the red region, facilitating imaging through thick, highly scattering tissues, and paving the way for advanced in vivo imaging applications.

Beyond imaging applications, our in-depth understanding of LumiFAST’s structural requirements and optimal fluorogen choices for efficient BRET enabled us to engineer novel intracellular biosensors. By introducing a protease cleavage site as a linker between pFAST and NanoLuc, we developed sensors that accurately report protease activity in living cells. Fine-tuning of the linker length was essential to maximize BRET efficiency while ensuring effective protease access to the cleavage site. Additionally, we leveraged BRET to monitor protein-protein interactions in living cells. Careful construct topology and linker optimization enabled to maximize BRET efficiency for optimal detection of interactions in living cells. Collectively, these advances lay the groundwork for the future design and in vivo application of sensitive bioluminescent biosensors.

## MATERIALS & METHODS

See supporting information

## DATA AVAILABILITY

Data supporting the findings of this study are available within the article and supplementary information, and are available from the corresponding authors upon request. The plasmids developed in this study may be requested from the corresponding author.

## ACKNOWLEDGMENTS

We thank the microscopy facility of the Institut de Biologie Paris Seine of Sorbonne University, and more particularly France Lam and Chloé Chaumeton for their assistance. This work has been supported by the Agence Nationale de la Recherche (ANR-21-CE42-0019-02 AmpliSens) and the Institut Universitaire de France.

## AUTHOR CONTRIBUTIONS

H.M and A.Ga. designed the overall project and wrote the paper. H.M and A.Ga. designed the experiments. H.M., A.Go., L.E.H. performed the experiments. Y.S. and L.J. synthesized HBP-2,5DM and HBR-2,5DOM. H.M. and A.Ga. analyzed the experiments.

## REFERENCES

(1) Love, A. C.; Prescher, J. A. Seeing (and Using) the Light: Recent Developments in Bioluminescence Technology. Cell Chem. Biol. 2020, 27 (8), 904–920.

(2) Syed, A. J.; Anderson, J. C. Applications of Bioluminescence in Biotechnology and Beyond. Chem. Soc. Rev. 2021, 50 (9), 5668–5705.

(3) Contag, C. H.; Spilman, S. D.; Contag, P. R.; Oshiro, M.; Eames, B.; Dennery, P.; Stevenson, D. K.; Benaron, D. A. Visualizing Gene Expression in Living Mammals Using a Bioluminescent Reporter. Photochem. Photobiol. 1997, 66 (4), 523–531.

(4) Shimomura, O.; Masugi, T.; Johnson, F.; Haneda, Y. Properties and Reaction Mechanism of the Bioluminescence System of the Deep-Sea Shrimp Oplophorus Gracilorostris. Biochemistry 1978, 17 (6), 994–998.

(5) Hall, M. P.; Unch, J.; Binkowski, B. F.; Valley, M. P.; Butler, B. L.; Wood, M. G.; Otto, P.; Zimmerman, K.; Vidugiris, G.; Machleidt, T.; Robers, M. B.; Benink, H. A.; Eggers, C. T.; Slater, M. R.; Meisenheimer, P. L.; Klaubert, D. H.; Fan, F.; Encell, L. P.; Wood, K. V. Engineered Luciferase Reporter from a Deep Sea Shrimp Utilizing a Novel Imidazopyrazinone Substrate. ACS Chem. Biol. 2012, 7 (11), 1848–1857.

(6) Suzuki, K.; Kimura, T.; Shinoda, H.; Bai, G.; Daniels, M. J.; Arai, Y.; Nakano, M.; Nagai, T. Five Colour Variants of Bright Luminescent Protein for Real-Time Multicolour Bioimaging. Nat. Commun. 2016, 7, 13718.

(7) Hattori, M.; Wazawa, T.; Orioka, M.; Hiruta, Y.; Nagai, T. Creating Coveted Bioluminescence Colors for Simultaneous Multi-Color Bioimaging. Sci. Adv. 2025, 11 (4), eadp4750.

(8) Chu, J.; Oh, Y.; Sens, A.; Ataie, N.; Dana, H.; Macklin, J. J.; Laviv, T.; Welf, E. S.; Dean, K. M.; Zhang, F.; Kim, B. B.; Tang, C. T.; Hu, M.; Baird, M. A.; Davidson, M. W.; Kay, M. A.; Fiolka, R.; Yasuda, R.; Kim, D. S.; Ng, H.-L.; Lin, M. Z. A Bright Cyan-Excitable Orange Fluorescent Protein Facilitates Dual-Emission Microscopy and Enhances Bioluminescence Imaging in Vivo. Nat. Biotechnol. 2016, 34 (7), 760–767.

(9) Hiblot, J.; Yu, Q.; Sabbadini, M. D. B.; Reymond, L.; Xue, L.; Schena, A.; Sallin, O.; Hill, N.; Griss, R.; Johnsson, K. Luciferases with Tunable Emission Wavelengths. Angew. Chem. Int. Ed Engl. 2017, 56 (46), 14556–14560.

(10) Benaissa, H.; Ounoughi, K.; Aujard, I.; Fischer, E.; Goïame, R.; Nguyen, J.; Tebo, A. G.; Li, C.; Le Saux, T.; Bertolin, G.; Tramier, M.; Danglot, L.; Pietrancosta, N.; Morin, X.; Jullien, L.; Gautier, A. Engineering of a Fluorescent Chemogenetic Reporter with Tunable Color for Advanced Live-Cell Imaging. Nat. Commun. 2021, 12 (1), 6989.

(11) Torrey, Z. R.; Halbers, L. P.; Scipioni, L.; Tedeschi, G.; Digman, M. A.; Prescher, J. A. A Versatile Bioluminescent Probe with Tunable Color. RSC Chem. Biol. 2024. 10.1039/d4cb00101j.

(12) Carrington, J. C.; Dougherty, W. G. A Viral Cleavage Site Cassette: Identification of Amino Acid Sequences Required for Tobacco Etch Virus Polyprotein Processing. Proc. Natl. Acad. Sci. U. S. A. 1988, 85 (10), 3391–3395.

(13) Kapust, R. B.; Waugh, D. S. Controlled Intracellular Processing of Fusion Proteins by TEV Protease. Protein Expr. Purif. 2000, 19 (2), 312–318.

(14) Sanchez, M. I.; Ting, A. Y. Directed Evolution Improves the Catalytic Efficiency of TEV Protease. Nat. Methods 2020, 17 (2), 167–174.

(15) Williams, D. J.; Puhl, H. L., 3rd; Ikeda, S. R. Rapid Modification of Proteins Using a Rapamycin-Inducible Tobacco Etch Virus Protease System. PLoS One 2009, 4 (10), e7474.

(16) Banaszynski, L. A.; Liu, C. W.; Wandless, T. J. Characterization of the FKBP-Rapamycin-FRB Ternary Complex. J. Am. Chem. Soc. 2005, 127, 4715–4721.

