## Supplementary Information for "Chemogenetic modulation of luciferase emission color for imaging and sensing"

**This pdf contains**

Supplementary Figures 1-11

Supplementary Table 1

Materials and Methods

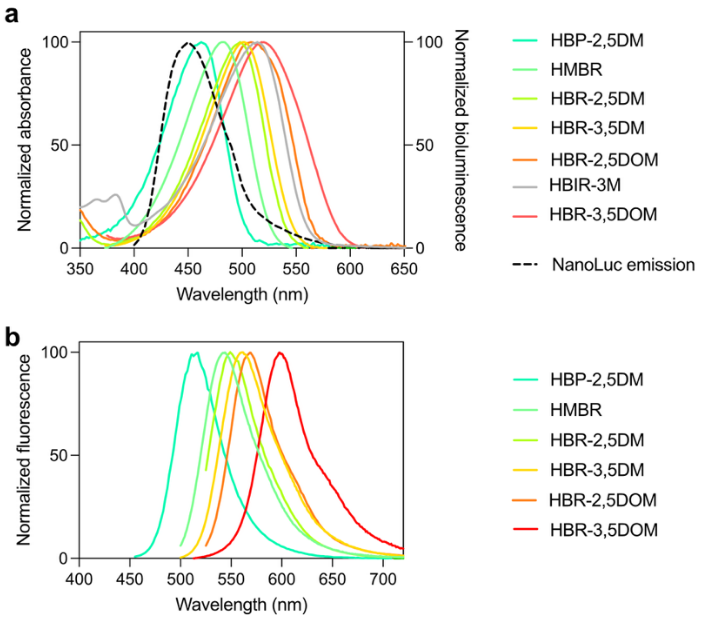


**Supplementary Figure 1. Spectral properties of pFAST with various fluorogens. a** Absorption spectra of various pFAST:fluorogen complexes in pH 7.4 PBS buffer. The emission spectra of NanoLuc is shown in dashed line. **b** Fluorescence emission spectra of various pFAST:fluorogen complex. **a,b** See also **Supplementary Table 1**. Data with HMBR, HBR-3,5DM, HBR-3,5DOM were previously reported in ref. 1**.**

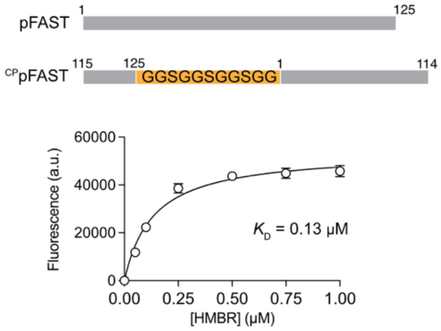


**Supplementary Figure 2. Characterization of $^{CP}$pFAST.** $^{CP}$pFAST was obtained by fusing the original N- and C-termini of pFAST with a GGSGGSGGSGG linker and introducing new N- and C- termini in position 115 and 114. The graph shows HMBR titration curves in pH 7.4 PBS buffer. The protein concentration was fixed to 0.05 µM. Data represent the mean ± SD of three experiments. Least-squares fit (line) gave the thermodynamic dissociation constant $K_D$ of the $^{CP}$pFAST:HMBR complex.

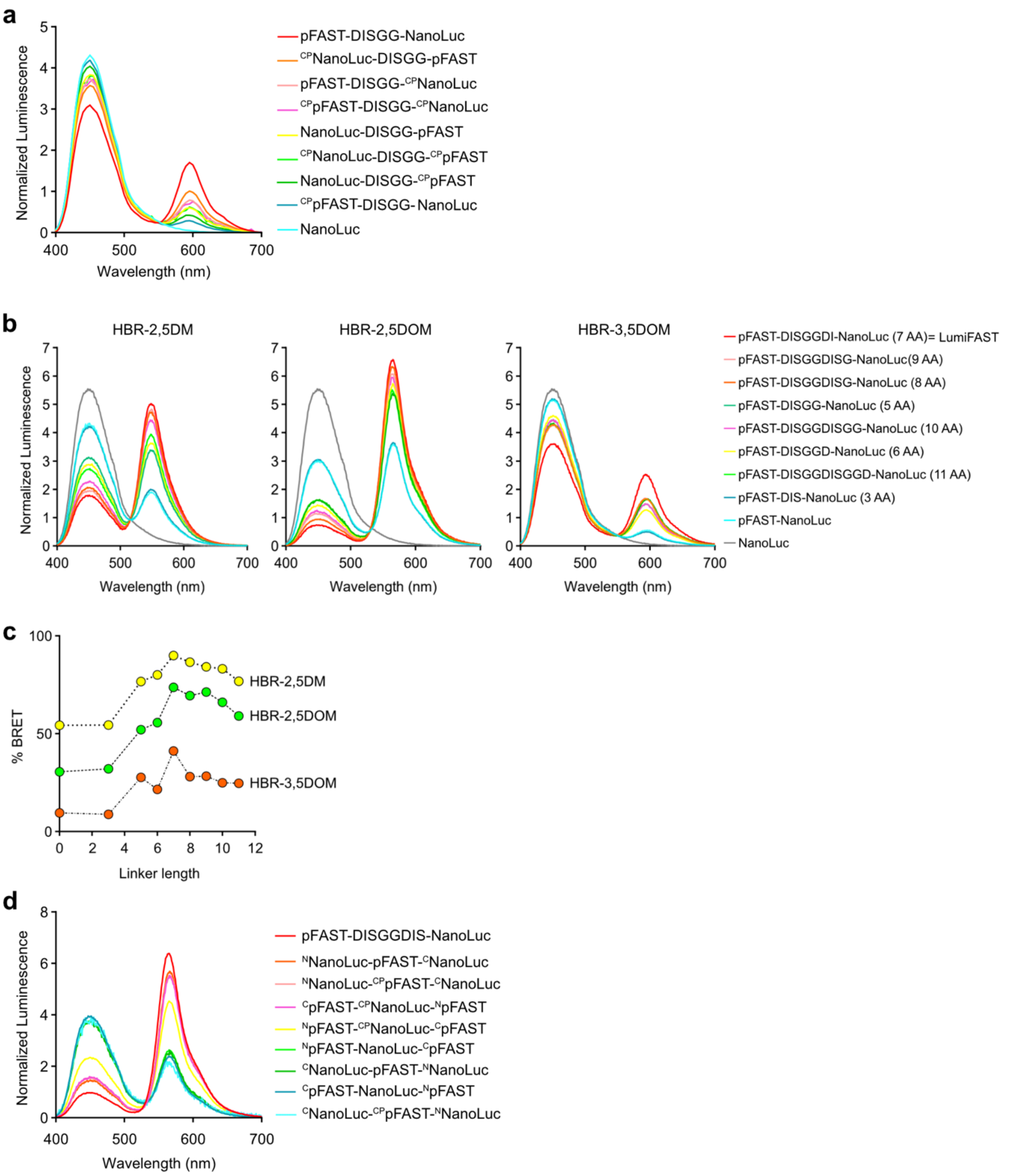


**Supplementary Figure 3. Engineering and optimization of LumiFAST. a** Bioluminescence emission spectra of various fusions of pFAST (or $^{CP}$pFAST) and NanoLuc (or $^{CP}$NanoLuc) (50 nM) with HBR-3,5DOM (5 $\mu$M) and with final 250-fold dilution of NanoGlo Furimazine substrate (Promega). **b** Bioluminescence emission spectra of various pFAST-linker-NanoLuc fusions (50 nM) with HBR-2,5DM, HBR-2,5DOM and HBR-3,5DOM (5 $\mu$M) and with final 250-fold dilution of NanoGlo Furimazine substrate (Promega). **c** Apparent BRET efficiency measured from the data shown in **b**. **d** Bioluminescence emission spectra of various insertions of pFAST (or $^{CP}$pFAST) into NanoLuc (or $^{CP}$NanoLuc) (50 nM) and vice versa with HBR-2,5DOM (5 $\mu$M) and with final 250-fold dilution of NanoGlo Furimazine substrate (Promega). **a,b,d** Spectra were normalized by the total bioluminescence intensity (area normalization).

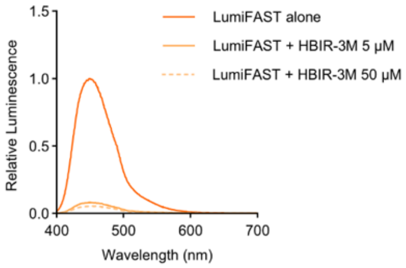


**Supplementary Figure 4. Bioluminescence of LumiFAST with the dark fluorogen HBIR-3M.** Bioluminescence emission spectra of LumiFAST alone or with HBIR-3M (at 5 μM or 50 μM) with final 250-fold dilution of NanoGlo Furimazine substrate (Promega). Spectra were normalized by the maximal bioluminescence intensity value of the condition LumiFAST alone.

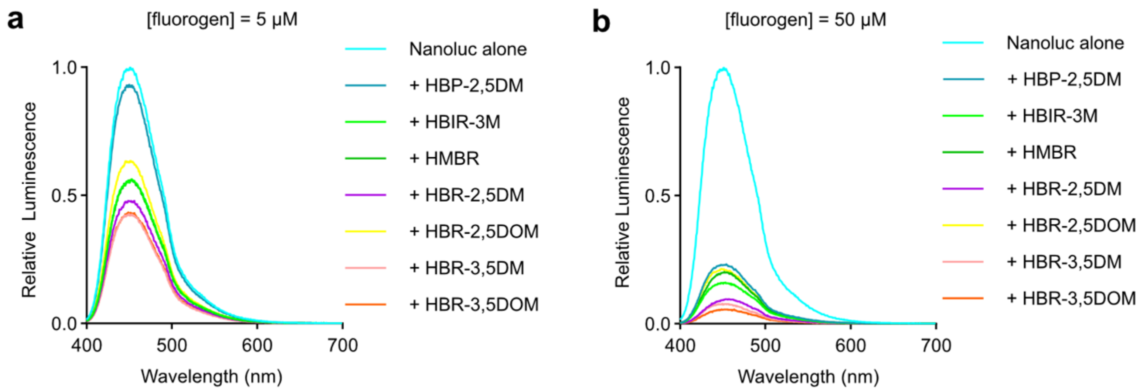


**Supplementary Figure 5. Bioluminescence of NanoLuc in the presence of free fluorogen.** Bioluminescence emission spectra of NanoLuc (50 nM) in presence of various fluorogens at 5 µM (**a**) or 50 µM (**b**) with final 250-fold dilution of NanoGlo Furimazine substrate (Promega). Spectra were normalized by the maximal bioluminescence intensity value of the condition NanoLuc alone.

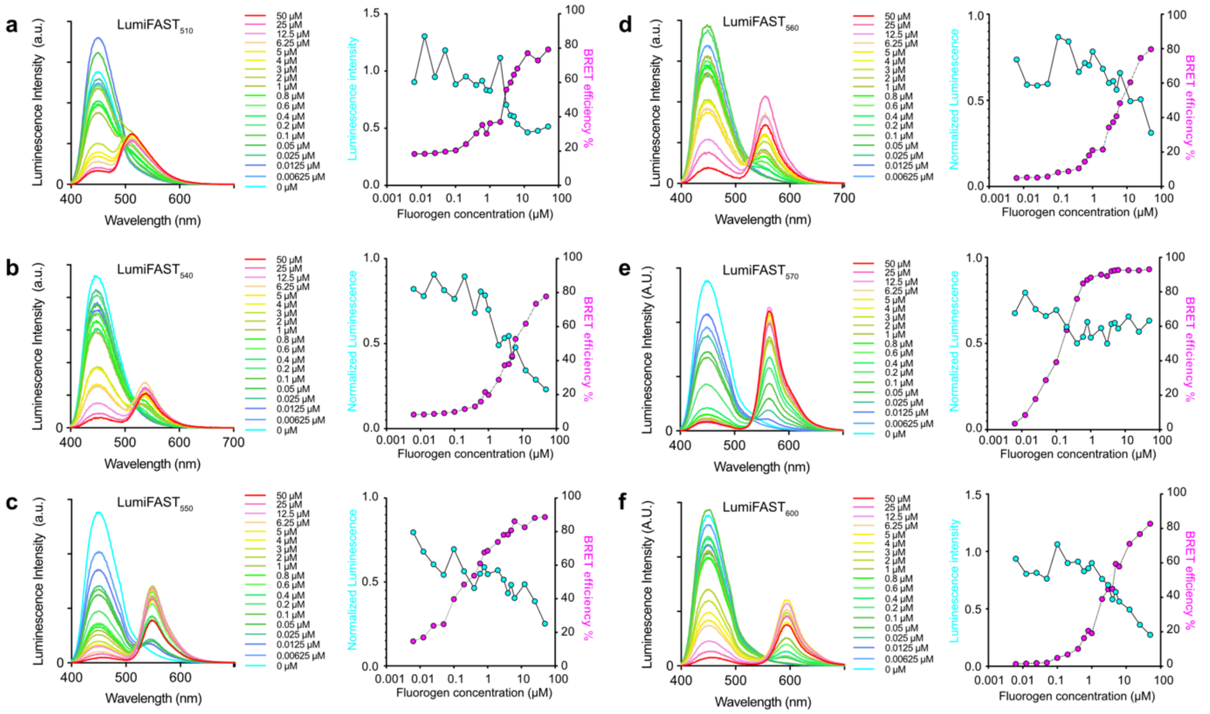


**Supplementary Figure 6. Bioluminescence of LumiFAST in function of fluorogen concentration.** Bioluminescence emission spectra of LumiFAST (50 nM) with various concentrations of HBP-2,5DM (**a**), HMBR (**b**), HBR-2,5DM (**c**), HBR-3,5DM (**d**), HBR-2,5DOM) (**e**) and HBR-3,5DOM (**f**) (with final 250-fold dilution of NanoGlo Furimazine substrate (Promega)). Graphs show the relative bioluminescence intensity and the apparent BRET efficiency at each fluorogen concentration.

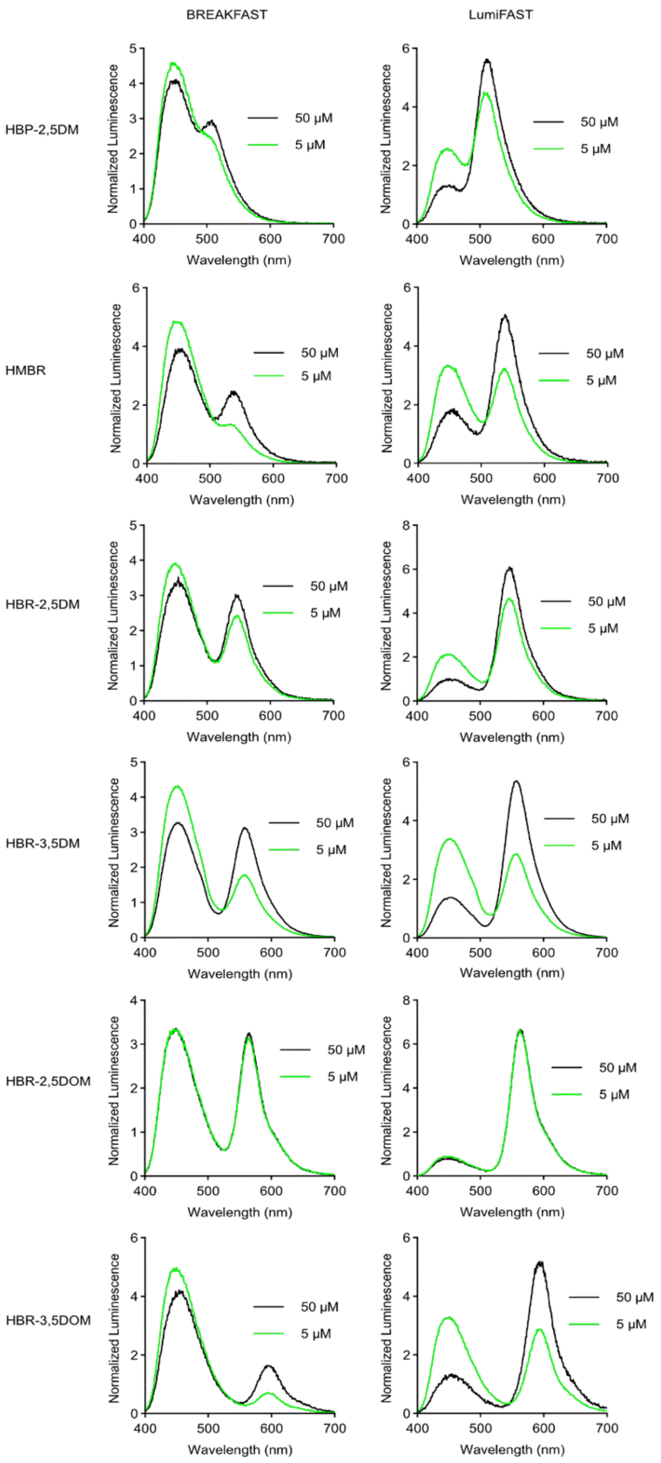


**Supplementary Figure 7. Comparison of LumiFAST and BREAKFAST**. Bioluminescence emission spectra of BREAKFAST and LumiFAST with various fluorogens at 5 or 50 μM (with final 250-fold dilution of NanoGlo Furimazine substrate (Promega)). Spectra were normalized by the total bioluminescence intensity (area normalization).

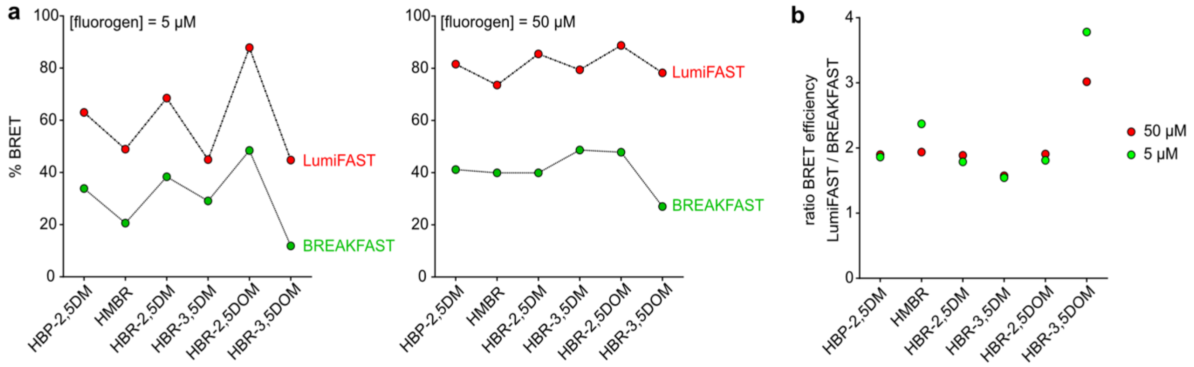


**Supplementary Figure 8. Comparison of LumiFAST and BREAKFAST**. **a** BRET efficiency of BREAKFAST and LumiFAST with various fluorogens at 5 or 50 µM (from data shown in **Supplementary Figure 6**). **b** Ratio of the BRET efficiency of LumiFAST and BREAKFAST.

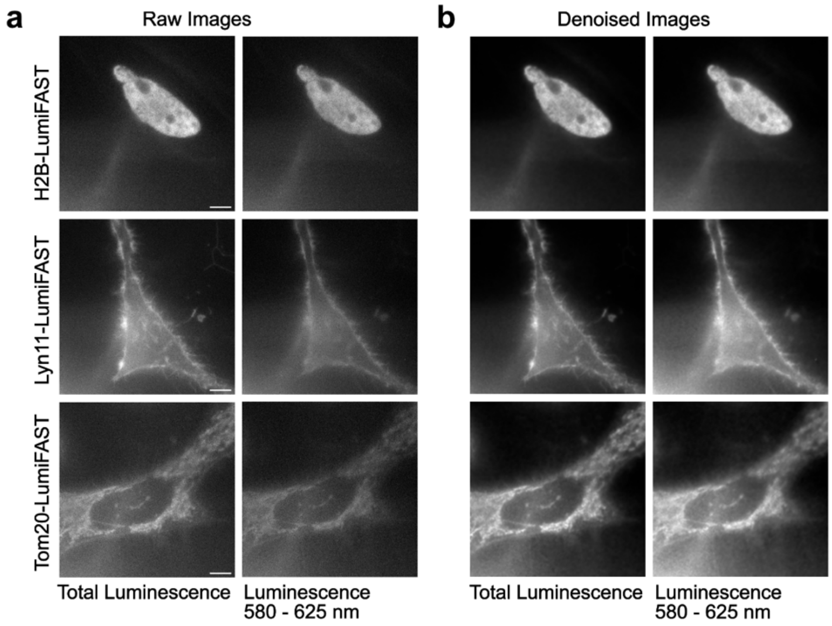


**Supplementary Figure 9. AI-assisted denoising of bioluminescence microscopy images. a** Raw bioluminescence images (see also **Figure 3**). **b** Denoised bioluminescence microscopy images obtained applying Noise2Void 2D deep-learning method, by training the network with high quality fluorescence microscopy images (see **Figure 3** for the fluorescence images).

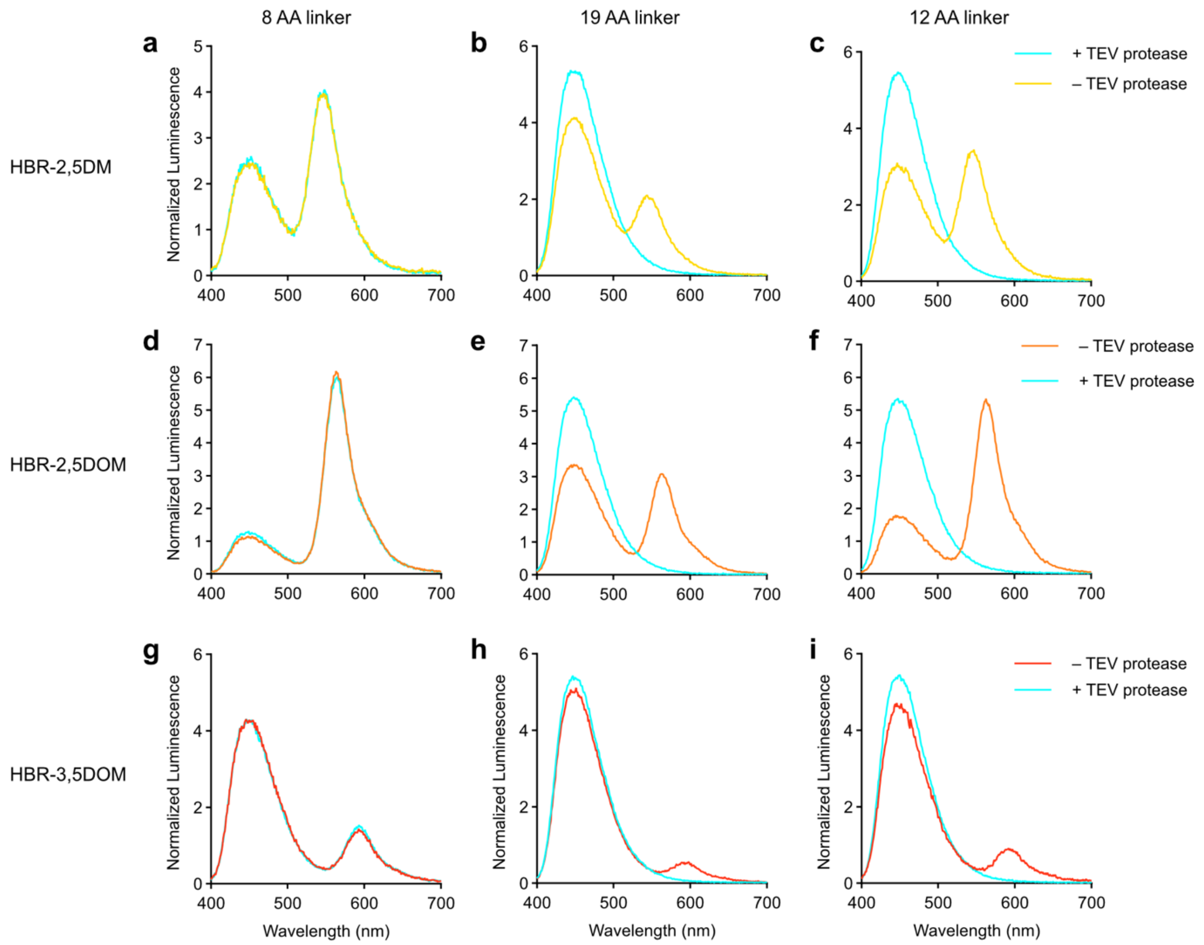


**Supplementary Figure 10. BRET-based sensing of protease activity.** Bioluminescence emission spectra of the three sensor candidates (50 nM) (fluorogen 5 μM with final 250 fold dilution of NanoGlo Furimazine substrate (Promega)) before and after treatment with the TEV protease. Spectra were normalized by the total bioluminescence intensity (area normalization). Sensors: **a,d,g** 8 AA linker ; **b,e,h** 19 AA linker ; **c,f,i** 12 AA linker. Fluorogens: **a-c** HBR-2,5DM ; **d-f** HBR-2,5DOM ; **g-i** HBR-3,5DOM.

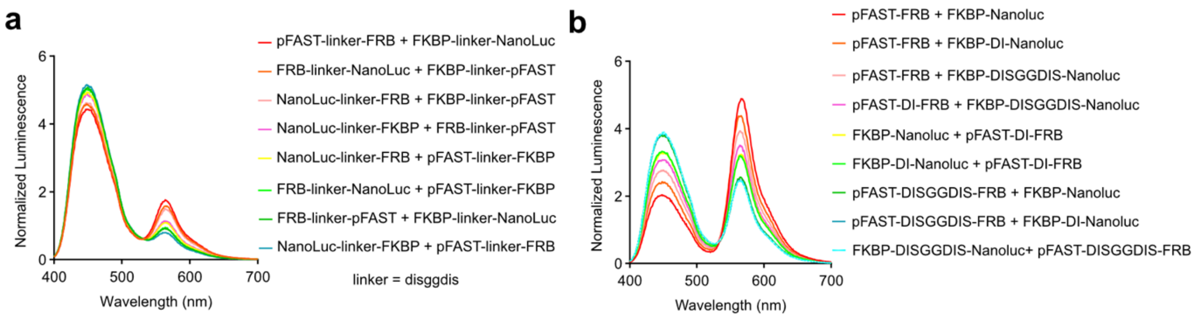


**Supplementary Figure 11. BRET-based sensing of protein-protein interactions. a** Bioluminescence emission spectra of solutions containing various FRB and FKBP fusions (50 nM each), HBR-2,5DOM (5 μM) and rapamycin (100 nM) with final 250-fold dilution of NanoGlo Furimazine substrate (Promega). Spectra were normalized by the total bioluminescence intensity (area normalization). **b** Influence of the linkers on the BRET efficiency. Bioluminescence emission spectra of solutions containing pFAST-linker-FRB (50 nM) and FKBP-linker-NanoLuc (25 nM), HBR-2,5DOM (5 μM), rapamycin (100 nM) with final 250-fold dilution of NanoGlo Furimazine substrate (Promega). Spectra were normalized by the total bioluminescence intensity (area normalization).

**Supplementary Table 1.** Properties of pFAST with various fluorogens in PBS pH 7.4

| fluorogen | $K_D$ (μM) | $\lambda_{abs}$ free fluorogen (nm) | $\lambda_{abs}$ (nm) | $\lambda_{em}$ (nm) | QY (%) | ε ($mM^{-1}cm^{-1}$) | Ref. |
|---|---|---|---|---|---|---|---|
| HBIR-3M | 0.04 | 426 | 514 | 567 | 0.3 | 12 | 1 |
| HBP-2,5DM | 0.044 | 368 | 462 | 517 | 18 | 41 | This study |
| HMBR | 0.01 | 401 | 481 | 542 | 23 | 54 | 1 |
| HBR-2,5DM | 0.044 | 406 | 498 | 549 | 28 | N.D. | This study |
| HBR-3,5DM | 0.01 | 402 | 501 | 561 | 44 | 49 | 1 |
| HBR-2,5DOM | 0.021 | 432 | 508 | 569 | 45 | 39 | This study |
| HBR-3,5DOM | 0.06 | 405 | 520 | 600 | 35 | 44 | 1 |

Abbreviations are as follows: $\lambda_{abs}$ wavelength of maximal absorption; $\lambda_{em}$ wavelength of maximal emission; ε, molar absorptivity at $\lambda_{abs}$; QY fluorescence quantum yield; $K_D$ thermodynamic dissociation constant.

### Materials and Methods

#### Organic synthesis

**General** Commercially available reagents were used as obtained. $^{1}H$ and $^{13}C$ NMR spectra were recorded at 300K on a Bruker AM 300 spectrometer; chemical shifts are reported in ppm with protonated solvent as internal reference; coupling constants *J* are given in Hz. The synthesis of HBIR-3M[1], HMBR[2], HBR-2,5DM[3], HBR-3,5DM[3], HBR-3,5DOM[3] was previously reported. These fluorogens are commercially available from the Twinkle Factory under the name $match_{dark}$, $^{TF}$Lime, $match_{550}$, $^{TF}$Amber, $^{TF}$Coral, respectively. The synthesis of HBR-2,5DOM was previously reported as well[4]. We present here an alternative protocol. We report also the synthesis of HBP-2,5DM.

**5-(4-hydroxy-2,5-dimethoxybenzylidene)-2-thioxothiazolidin-4-one (HBR-2,5DOM)**
2,5-dimethoxy-4-hydroxybenzaldehyde (100 mg, 0.549 mmol) and rhodanine (73 mg, 0.549 mmol) powders were mixed in a test tube and heated to 170°C. The solids quickly melt then re-solidify after 5 min. After cooling down, the solid residue was dissolved in 35 mL of hot (70°C) ethanol. Addition of 70 mL of water caused an orange precipitate to form, which was filtered and washed thrice with water. After drying over $P_2O_5$, 101 mg of **HBR-2,5DOM** (62% yield) was obtained as an orange powder. $^{1}H$ NMR (300 MHz, DMSO) $\delta$ 13.62 (s, 1H), 10.31 (s, 1H), 7.76 (s, 1H), 6.87 (s, 1H), 6.6 (s, 1H) 2, 3.82 (s, 3H), 3.80 (s, 3H). $^{13}C$ NMR (75 MHz, DMSO) $\delta$ 195.55, 169.57, 154.85, 152.29, 142.32, 127.65, 120.18, 112.92, 111.61, 100.48, 56.19, 55.97. HRMS (ESI) m/z 296.0052 $[M–H]^-$ (calculated for $[C_{12}H_{10}NO_4S_2]^-$ 296.0051)

**5-(4-hydroxy-2,5-dimethylbenzylidene)-2-iminothiazolidin-4-one (HBP-2,5DM)**
2,5-dimethyl-4-hydroxybenzaldehyde (155 mg, 1.033 mmol) and pseudothiohydantoin (100 mg, 0.861 mmol) powders were mixed in a test tube and heated to 170°C. The solids quickly melt then re-solidify after 5 min. After cooling down, the solid residue was dissolved in 7 mL of hot (70°C) ethanol. Addition of 15 mL of water caused a dark red precipitate to form, which was filtered and washed thrice with water. The precipitate was filtered and purified by repeating the workup protocol (dissolve in 7 mL hot ethanol, add 15 mL of water, filter the product and wash it with water). After drying over $P_2O_5$, 69 mg of **HBP-2,5DM** (32% yield) was obtained as a red powder. $^{1}H$ NMR (300 MHz, DMSO) $\delta$ 9.86 (s, 1H), 9.26 (s, 1H), 8.96 (s, 1H), 7.63 (s, 1H), 7.19 (s, 1H), 6.71 (s, 1H), 2.28 (s, 3H), 2.12 (s, 3H). $^{13}C$ NMR (75 MHz, DMSO) $\delta$ 180.42, 175.78, 156.82, 137.72, 129.53, 126.66, 126.58, 123.57, 122.02, 116.99, 19.17, 15.70. HRMS (ESI) m/z 249.0690 $[M+H]^+$ (calculated for $[C_{12}H_{13}N_2O_2S]^+$ 249.0692)

### Biology

#### General

PCR reactions were performed with Q5 polymerase (New England Biolabs) in the buffer provided. PCR products were purified using QIAquick PCR purification kit (Qiagen). DNAse I, T4 ligase, Fusion polymerase, Taq ligase and Taq exonuclease were purchased from New England Biolabs and used with accompanying buffers and according to manufacturer protocols. Isothermal assemblies (Gibson assembly) were performed using homemade mix prepared according to previously described protocols[5]. Small-scale isolation of plasmid DNA was done using QIAprep miniprep kit (Qiagen) from 4 mL overnight bacterial culture supplemented with appropriate antibiotics. Large-scale isolation of plasmid DNA was done using the QIAprep maxiprep kit (Qiagen) from 150 mL of overnight bacterial culture supplemented with appropriate antibiotics. All plasmid sequences were confirmed by Sanger sequencing with appropriate sequencing primers (GATC Biotech). All the plasmids used in this study are listed, as well as proteins sequences.

#### Cloning

All E. coli expression plasmids were constructed using the pET28 backbone. The plasmids have been generated using isothermal Gibson assembly or restriction enzymes and allow protein expression of relative insert. For isothermal Gibson assembly, two linear DNA fragments of the pET28 vector were generated by PCR. The plasmid pAG952 allowing the bacterial expression of $^{CP}$pFAST was construct by Gibson assembly from the plasmid pAG882[6]. The plasmid pAG1481 for the bacterial expression of NanoLuc was generated by inserting the amplified NanoLuc sequence from the plasmid 87696 from Promega in a pET28 empty vector using the BamHI and NotI restriction sites. The plasmids pAG1482 and pAG1485 for the bacterial expression of NanoLuc-DISGG-pFAST and pFAST-DISGG-NanoLuc were generated by inserting the NanoLuc-DISGG-pFAST and pFAST-DISGG-NanoLuc sequences respectively in the empty pET28 vector. The sequence of $^{CP}$NanoLuc was previously described by Johnsson et al.[7]. They have fused the N-terminal and the C-terminal with a linker (GGTGGS)*2 and further created new extremities between the amino acids 65 and 66. The plasmid pAG1493 for the bacterial expression of $^{CP}$NanoLuc was generated by inserting the amplified $^{CP}$NanoLuc sequence in the pET28 vector with restriction enzymes. The plasmids pAG1486 and pAG1487 for the bacterial expression of $^{CP}$NanoLuc-DISGG-pFAST and pFAST-DISGG-$^{CP}$NanoLuc were generated by inserting the amplified $^{CP}$NanoLuc-pFAST and pFAST-$^{CP}$NanoLuc sequences respectively in the empty pET28 vector. The $^{CP}$pFAST sequence was obtained from the plasmid pAG952. The plasmids pAG1489 and pAG1490 for the bacterial expression of $^{CP}$pFAST-DISGG-NanoLuc and NanoLuc-$^{CP}$pFAST were generated by inserting the amplified $^{CP}$pFAST-NanoLuc and NanoLuc-$^{CP}$pFAST sequences respectively in the empty pET28 vector. The plasmids pAG1491 and pAG1492 for the bacterial expression of $^{CP}$NanoLuc-$^{CP}$pFAST and $^{CP}$pFAST-$^{CP}$NanoLuc were generated by inserting the amplified $^{CP}$NanoLuc-$^{CP}$pFAST and $^{CP}$pFAST-$^{CP}$NanoLuc respectively in the empty pET28 vector. All the plasmids expressing pFAST-NanoLuc with different sizes of the linker (pAG 1546,1547,1548,1549, 1621,1622,1623 and 1623) were generated by Gibson of 2 PCR fragments amplified of the plasmid pAG1485. The plasmids used for the insertion strategy were generated by Gibson. The plasmid pAG1566 for the bacterial expression of $^{C}$pFAST-$^{CP}$NanoLuc-$^{N}$pFAST was generated by inserting the $^{CP}$NanoLuc sequence in the sequence of $^{CP}$pFAST in the plasmid pAG952. The plasmid pAG1567 for the bacterial expression of $^{C}$pFAST-NanoLuc-$^{N}$pFAST was generated by inserting the NanoLuc sequence in the sequence of the $^{CP}$pFAST of the plasmid pAG952. The plasmid pAG1568 for the bacterial expression of $^{C}$NanoLuc-$^{CP}$pFAST-$^{N}$NanoLuc was generated by inserting the $^{CP}$pFAST sequence in the sequence of the $^{CP}$NanoLuc in the plasmid pAG1493. The plasmid pAG1569 for the bacterial expression of $^{C}$NanoLuc-pFAST-$^{N}$NanoLuc was generated by inserting the pFAST sequence in the sequence of the $^{CP}$NanoLuc in the plasmid pAG1493. The plasmid pAG1570 for the bacterial expression of $^{N}$NanoLuc-$^{CP}$pFAST-$^{C}$NanoLuc was generated by inserting the $^{CP}$pFAST sequence in the sequence of the NanoLuc in the plasmid pAG1481. The plasmid pAG1571 for the bacterial expression of $^{N}$NanoLuc-pFAST-$^{C}$NanoLuc was generated by inserting the pFAST sequence in the sequence of the NanoLuc of the plasmid pAG1481. The plasmid pAG1572 for the bacterial expression of $^{N}$pFAST-$^{CP}$NanoLuc-$^{C}$pFAST was generated by inserting the $^{CP}$NanoLuc sequence in the sequence of the pFAST in the plasmid pAG641. The plasmid pAG1573 for the bacterial expression of $^{N}$pFAST-NanoLuc-$^{C}$pFAST was generated by inserting the NanoLuc sequence in the sequence of pFAST in the plasmid pAG641.

The plasmid pAG1654 for mammalian expression of LumiFAST was generated by replacing the reported sequence of nirFAST in the plasmid pAG1375[8] with the sequence of LumiFAST. The plasmid pAG1790 for mammalian expression of H2B-LumiFAST was generated by inserting the sequence of H2B at the N-terminal of LumiFAST in the plasmid pAG1654. The plasmid pAG1802 for mammalian expression of Tom20-LumiFAST was generated by inserting the sequence of Tom20 at the N-terminal of LumiFAST in the plasmid pAG1654. The plasmid pAG1857 for mammalian expression of Lyn11-LumiFAST was generated by inserting the sequence of Lyn11 at the N-terminal of LumiFAST in the plasmid pAG1654. The plasmid pAG1798 for mammalian expression of H2B-NanoLuc-iRFP670 was generated by Gibson from the plasmids pAG1436 and pAG1420. The plasmid pAG1799 expressing Tom20-NanoLuc-iRFP670 was generated by Gibson from the plasmids pAG1802 and pAG1420.

The plasmid pAG1625 for the bacterial expression of pFAST-DENLYFQS-NanoLuc was generated from pAG1622 by including the cleavage site using primers with overhangs encoding the cleavage site. The plasmid pAG1670 for the bacterial expression of pFAST-DISGGDENLYFQSDISGGD-NanoLuc was generated from the plasmid pAG1625 by generating two DNA fragments with overhangs encoding for the cleavage site. The plasmid pAG1797 for the bacterial expression of pFAST-DISENLYFQSDI-NanoLuc was generated from the plasmid pAG1625 by generating two DNA fragments with overhangs encoding for the cleavage site. The plasmid pAG1656 for mammalian expression of the TEV-p sensor was generated by inserting the TEV cleavage site sequence into the LumiFAST sequence in the plasmid pAG1654.Two DNA fragments were generated by PCR with overhangs encoding for the cleavage site.

The plasmids used for the protein-protein interactions study were generated by Gibson strategy. The plasmid pAG1601 for the bacterial expression of FRB-DISGGDIS-NanoLuc was generated by inserting the FRB sequence at the N-terminal of the NanoLuc sequence of the plasmid pAG1548. The plasmid pAG1602 for the bacterial expression of FRB-DISGGDIS-pFAST was generated by inserting the FRB sequence at the N-terminal of the pFAST sequence of the plasmid pAG1486. The plasmid pAG1603 for the bacterial expression of FKBP-DISGGDIS-NanoLuc was generated by inserting the FKBP sequence at the N-terminal of the NanoLuc sequence of the plasmid pAG1548. The plasmid pAG1604 for the bacterial expression of FKBP-DISGGDIS-pFAST was generated by inserting the FKBP sequence at the N-terminal of the pFAST sequence of the plasmid pAG1486. The plasmid pAG1703 for the bacterial expression of NanoLuc-DISGGDIS-FRB was generated by inserting the FRB sequence at the C-terminal of the NanoLuc sequence of the plasmid pAG1481. The plasmid pAG1704 for the bacterial expression of pFAST-DISGGDIS-FRB was generated by inserting the FRB sequence at the C-terminal of the pFAST sequence of the plasmid pAG1622. The plasmid pAG1705 for the bacterial expression of NanoLuc-DISGGDIS-FKBP was generated by inserting the FKBP sequence at the C-terminal of the NanoLuc sequence of the plasmid pGA1481. The plasmid pAG1706 for the bacterial expression of pFAST-DISGGDIS-FKBP was generated by inserting FKBP sequence at C-terminal of the pFAST sequence of the plasmid pAG1622. The plasmids pAG1657 and pAG1658 for the bacterial expression of pFAST-FRB with different size of the linker were generated by Gibson of two PCR fragments amplified of the plasmid pAG1704. The plasmids pAG1659 and pAG1660 for bacterial expression of FKBP-NanoLuc with different size of the linker were generated by Gibson of two PCR fragments amplified of the plasmid pAG1603. The plasmid pAG1821 for mammalian of pFAST-FRB was generated by inserting the pFAST-FRB sequence in a CMV vector. The plasmid pAG1822 for mammalian of FKBP-NanoLuc was generated by inserting the FKBP-NanoLuc sequence in a CMV vector.

**Protein Expression, Purification, and *in vitro* experiments.**

Expression vectors of all proteins with an N-terminal His-tag under the control of a T7 promoter were transformed in BL21 *Escherichia coli* competent cells. Bacterial cells were grown at 37 °C in Lysogeny Broth (LB) supplemented with kanamycin (50 μg/mL) until $OD_{600nm}$ = 0.6. Expression was induced overnight at 16°C by the addition of isopropyl β-D-1-thiogalactopyranoside (IPTG) (1 mM). Cells were harvested by centrifugation (4000×g for 20 min at 4°C) and stored at –30°C. The cell pellets were resuspended in lysis buffer (PBS supplemented with 2.5 mM $MgCl_2$, protease inhibitor PMSF 1 mM, DNase 0.025 mg/mL) and sonicated (5 min, 20% of amplitude) on ice. The lysate was incubated for 2 h on ice to allow DNA digestion by DNase. Cellular fragments were removed by centrifugation (9,000×g for 1 h at 4 °C). The supernatant was incubated overnight at 4 °C by gentle agitation with pre-washed Co-NTA agarose beads in PBS buffer complemented with 10 mM of imidazole. Beads were washed with ten volumes of PBS complemented with 10 mM of imidazole. His-tagged proteins were eluted with five volumes of PBS complemented with 150 mM of imidazole. The buffer was exchanged to PBS (0.05 M phosphate buffer and 0.150 M NaCl) or HEPES buffer (HEPES 50 mM, NaCl 259 mM, pH 7.4) for calcium sensors using PD-10 desalting columns. The purity of the proteins was evaluated using SDS–PAGE electrophoresis stained with Coomassie blue.

**Physicochemical measurements**

Steady-state UV-Vis and fluorescence spectra were recorded at 25 °C on a Spark 10 M (Tecan). Data were processed using GraphPad Prism v.10.0.3. Fluorescence quantum yields, molar absorption coefficients and thermodynamic dissociation constants were determined as previously described[8].

**Bioluminescence measurements**

Proteins were diluted to 50 nM and reactions were performed with a final 250-fold dilution of NanoGlo furimazine substrate (Promega). Experiments were performed using a 1 cm long cuvette in 100 μL volume at RT. Bioluminescence spectra were measured using a Jasco FP-8300 spectrofluorometer (Jasco Inc., Easton, MD, USA) with 1 nm emission bandwidth, high sensitivity and a scanning speed of 500 nm/min, and blocking the excitation to record only the bioluminescence light. BRET efficiencies were evaluated by computing:

$$\text{BRET efficiency} = I_{\text{max, acceptor}} / (I_{\text{max, donor}} + I_{\text{max, acceptor}})$$

Where $I_{max, acceptor}$ is the maximal intensity of the acceptor band and $I_{max, donor}$ is the maximal intensity of the donor band.

**Thermodynamic analysis**

The $EC_{50}$ for rapamycin was determined using 5 μM of the fluorogen HBR-2,5DOM, 50 nM of the pFAST-FRB protein, 25 nM of the FKBP-NanoLuc protein, a final dilution of 250-fold of Furimazine and various concentrations of rapamycin. The luminescence intensity of the BRET-based protein-protein interactions was plotted as a function of Rapamycin concentration.

**Cell culture**

HeLa cells (ATCC CRM-CCL2) were cultured in Minimal Essential Media (MEM) supplemented with phenol red, Glutamax I, 1 mM of sodium pyruvate, 1% (vol/vol) of non-essential amino-acids and 10% (vol/vol) fetal calf serum (FCS), at 37 °C in a 5% $CO_2$ atmosphere. For imaging, cells were seeded in μDish IBIDI (Biovalley) coated with poly-L-lysine. Transient transfections with 1 μg of the appropriate plasmids were performed using Genejuice (Merck) according to the manufacturer's protocols for 24 h prior to imaging. Cells were washed with DPBS (Dulbecco's Phosphate-Buffered Saline) and treated with DMEM media (without serum and phenol red) supplemented with the compounds at the indicated concentration.

**Bioluminescence imaging, micrographs denoising and data analysis**

The bioluminescent micrographs were acquired on a Nikon TIRF microscope motorized stage XY and perfect focus system (PFS) with an iXon EMCCD camera (Andor) integrated in Metamorph software by Gataca Systems. Images for luciferase activity were obtained using NanoGlo substrate diluted 50 times. For each single micrograph, a fluorescent image was generated before adding NanoLuc substrate. During the bioluminescence imaging, the laser excitation of the microscope was shut down. The whole emitted light was acquired over 10 s without emission filters; specific fluorogens signals were acquired over 10 s using the following emission filters: 420-480 nm, 500-550 nm, 580-625 nm and 670-780 nm. The images were analyzed using Fiji (Image J) and NIS-Elements (Nikon). Images **Supplementary Figure 9**, **Figure 6d** and **Figure 7d** were enhanced by Noise2Void on Fiji (Image J). The FIJI neural network was trained using a fluorescent micrograph. The signal to noise ratio was improved using the trained neural network.

**Phantom tissue generation**

The phantom tissue was generated to replicate brain grey optical properties. The components were in these following ratios: 2 g of agar and 300 mg of aluminum oxide mixed in an Erlenmeyer containing 100 mL de-ionized water and a magnetic stir bar. The mixture was heated while stirring until the temperature reached 94°C. Once this temperature reached, the mixture was moved to a separate stir plate at 55°C to prevent early firming and 300 µL India ink was added and stirred until completely mixed.

The mixture was then poured in Petri dishes for molding. 10 mm was the depth chosen and marked on the Petri dishes beforehand. This allowed us to have very precise thickness for the phantom system. The dishes were then placed at 4°C to rapidly solidify to prevent precipitation of the aluminum oxide and then were left at 4°C for at least 24 h.

**Protein and DNA sequences:**

pFAST, Nanoluc, $^{cp}$pFAST, $^{cp}$Nanoluc

LumiFAST :

MEHVAFGSEDIENTLANMDDEQLDRLAFGVIQLDGDGNILLYNAAEGDITGRDPKQVIGKNF
FKDVAPGTDTPEFYGKFKEGAASGNLNTMFEWTIPTSRGPTKVKVHLKKALSGDRYWVFV
KRVDISGGDIVFTLEDFVGDWRQTAGYNLDQVLEQGGVSSLFQNLGVSVTPIQRIVLSGEN
GLKIDIHVIIPYEGLSGDQMGQIEKIFKVVYPVDDHHFKVILHYGTLVIDGVTPNMIDYFGRPY
EGIAVFDGKKITVTGTLWNGNKIIDERLINPDGSLLFRVTINGVTGWRLCERILA

$^{CP}$pFAST:
MGDRYWVFVKRVGGSGGSGGSGGEHVAFGSEDIENTLANMDDEQLDRLAFGVIQLDGDG
NILLYNAAEGDITGRDPKQVIGKNFFKDVAPGTDTPEFYGKFKEGAASGNLNTMFEWTIPTS
RGPTKVKVHLKKALS

$^{CP}$Nanoluc:
MGLSGDQMGQIEKIFKVVYPVDDHHFKVILHYGTLVIDGVTPNMIDYFGRPYEGIAVFDGKKI
TVTGTLWNGNKIIDERLINPDGSLLFRVTINGVTGWRLCERILAGGTGGSGGTGGSMVFTLE
DFVGDWRQTAGYNLDQVLEQGGVSSLFQNLGVSVTPIQRIVLSGENGLKIDIHVIIPYE

Tom20:
VGRNSAIAAGVCGALFIGYCIYFDRKRRSDPNF

Lyn11:
GCIKSKGKDSA

H2B:
PEPAKSAPAPKKGSKKAVAKTQKKGDKKRRKTRKESYAIYVYKVLKQVHPDTGISSKAMGI
MNSFVNDIFERIAGEASRLAHYNKRSTITSREIQTAVRLLLPGELAKHAVSEGTKAVTKYTSS
K

TEV cleavage site:
ENLYFQS

FRB:

MILWHEMWHEGLEEASRLYFGERNVKGMFEVLEPLHAMMERGPQTLKETSFNQAYGRDL
MEAQEWCRKYMKSGNVKDLLQAWDLYYHVFRRISK

FKBP:

MGVQVETISPGDGRTFPKRGQTCVVHYTGMLEDGKKFDSSRDRNKPFKFMLGKQEVIRG
WEEGVAQMSVGQRAKLTISPDYAYGATGHPGIIPPHATLVFDVELLKLEE

| Plasmid number | Vector | ORF | Sequence |
|---|---|---|---|
| 1481 | pET28 | NanoLuc | atggtcttcacactcgaagatttcgttggggactggcgacagacagccggctacaacctggaccaagtccttga<br>acagggaggtgtgtccagtttgtttcagaatctcggggtgtccgtaactccgatccaaaggattgtcctgagcggt<br>gaaaatgggctgaagatcgacatccatgtcatcatcccgtatgaaggtctgagcggcgaccaaatgggccag<br>atcgaaaaaatttttaaggtggtgtaccctgtggatgatcatcactttaaggtgatcctgcactatggcacactggt<br>aatcgacggggttacgccgaacatgatcgactatttcggacggccgtatgaaggcatcgccgtgttcgacggc<br>aaaaagatcactgtaacagggaccctgtggaacggcaacaaaattatcgacgagcgcctgatcaaccccga<br>cggctccctgctgttccgagtaaccatcaacggagtgaccggctggcggctgtgcgaacgcattctggcg |
| 1482 | pET28 | NanoLuc-DISGG-pFAST | atggtcttcacactcgaagatttcgttggggactggcgacagacagccggctacaacctggaccaagtccttga<br>acagggaggtgtgtccagtttgtttcagaatctcggggtgtccgtaactccgatccaaaggattgtcctgagcggt<br>gaaaatgggctgaagatcgacatccatgtcatcatcccgtatgaaggtctgagcggcgaccaaatgggccag<br>atcgaaaaaatttttaaggtggtgtaccctgtggatgatcatcactttaaggtgatcctgcactatggcacactggt<br>aatcgacggggttacgccgaacatgatcgactatttcggacggccgtatgaaggcatcgccgtgttcgacggc<br>aaaaagatcactgtaacagggaccctgtggaacggcaacaaaattatcgacgagcgcctgatcaaccccga<br>cggctccctgctgttccgagtaaccatcaacggagtgaccggctggcggctgtgcgaacgcattctggcggat<br>attagcggcggcatggagcatgttgcctttggcagtgaggacatcgagaacactctggccaatatggacgacg<br>aacaactggataggttggcctttggcgtaattcagctcgatggtgacgggaatatcctgctgtacaatgctgctga<br>aggggacatcactggcagagatcccaaacaggtgattgggaagaacttcttcaaggatgttgcacctggaac<br>ggatactcccgagttttacggcaaattcaaggaaggcgcagcgtcagggaatctgaacaccatgttcgaatgg<br>acgataccgacaagcaggggaccaaccaaggtcaaggtgcacttgaagaaagccctttccggtgacagat<br>attgggtctttgtgaaacgggtgtaa |
| 1485 | pET28 | pFAST-DISGG-NanoLuc | atggagcatgttgcctttggcagtgaggacatcgagaacactctggccaatatggacgacgaacaactggata<br>ggttggcctttggcgtaattcagctcgatggtgacgggaatatcctgctgtacaatgctgctgaaggggacatca<br>ctggcagagatcccaaacaggtgattgggaagaacttcttcaaggatgttgcacctggaacggatactcccga<br>gttttacggcaaattcaaggaaggcgcagcgtcagggaatctgaacaccatgttcgaatggacgataccgac<br>aagcaggggaccaaccaaggtcaaggtgcacttgaagaaagccctttccggtgacagatattgggtctttgtg<br>aaacgggtggatattagcggcggcgtcttcacactcgaagatttcgttggggactggcgacagacagccggct<br>acaacctggaccaagtccttgaacagggaggtgtgtccagtttgtttcagaatctcggggtgtccgtaactccga<br>tccaaaggattgtcctgagcggtgaaaatgggctgaagatcgacatccatgtcatcatcccgtatgaaggtctg<br>agcggcgaccaaatgggccagatcgaaaaaatttttaaggtggtgtaccctgtggatgatcatcactttaaggt<br>gatcctgcactatggcacactggtaatcgacggggttacgccgaacatgatcgactatttcggacggccgtatg<br>aaggcatcgccgtgttcgacggcaaaaagatcactgtaacagggaccctgtggaacggcaacaaaattatc<br>gacgagcgcctgatcaaccccgacggctccctgctgttccgagtaaccatcaacggagtgaccggctggcgg<br>ctgtgcgaacgcattctggcg |
| 1486 | pET28 | $^{CP}$Nanoluc-DISGG-pFAST | ggtctgagcggcgaccaaatgggccagatcgaaaaaatttttaaggtggtgtaccctgtggatgatcatcacttt<br>aaggtgatcctgcactatggcacactggtaatcgacggggttacgccgaacatgatcgactatttcggacggcc<br>gtatgaaggcatcgccgtgttcgacggcaaaaagatcactgtaacagggaccctgtggaacggcaacaaaa<br>ttatcgacgagcgcctgatcaaccccgacggctccctgctgttccgagtaaccatcaacggagtgaccggctg<br>gcggctgtgcgaacgcattctggcgggcggcaccggcggcagcggcggcaccggcggcagcatggtcttc<br>acactcgaagatttcgttggggactggcgacagacagccggctacaacctggaccaagtccttgaacaggga<br>ggtgtgtccagtttgtttcagaatctcggggtgtccgtaactccgatccaaaggattgtcctgagcggtgaaaatg<br>ggctgaagatcgacatccatgtcatcatcccgtatgaagatattagcggcggcATGGAGCATGTTGC<br>CTTTGGCAGTGAGGACATCGAGAACACTCTGGCCAATATGGACGACGAAC<br>AACTGGATAGGTTGGCCTTTGGCGTAATTCAGCTCGATGGTGACGGGAATA<br>TCCTGCTGTACAATGCTGCTGAAGGGGACATCACTGGCAGAGATCCCAAA<br>CAGGTGATTGGGAAGAACTTCTTCAAGGATGTTGCACCTGGAACGGATACT<br>CCCGAGTTTTACGGCAAATTCAAGGAAGGCGCAGCGTCAGGGAATCTGAA<br>CACCATGTTCGAATGGACGATACCGACAAGCAGGGGACCAACCAAGGTCA<br>AGGTGCACTTGAAGAAAGCCCTTTCCGGTGACAGATATTGGGTCTTTGTGA<br>AACGGGTG |
| 1487 | pET28 | pFAST-DISGG-$^{CP}$Nanoluc | ATGGAGCATGTTGCCTTTGGCAGTGAGGACATCGAGAACACTCTGGCCAAT<br>ATGGACGACGAACAACTGGATAGGTTGGCCTTTGGCGTAATTCAGCTCGAT<br>GGTGACGGGAATATCCTGCTGTACAATGCTGCTGAAGGGGACATCACTGG<br>CAGAGATCCCAAACAGGTGATTGGGAAGAACTTCTTCAAGGATGTTGCACC<br>TGGAACGGATACTCCCGAGTTTTACGGCAAATTCAAGGAAGGCGCAGCGT<br>CAGGGAATCTGAACACCATGTTCGAATGGACGATACCGACAAGCAGGGGA<br>CCAACCAAGGTCAAGGTGCACTTGAAGAAAGCCCTTTCCGGTGACAGATAT<br>TGGGTCTTTGTGAAACGGGTGgatattagcggcggcggtctgagcggcgaccaaatgggcca<br>gatcgaaaaaatttttaaggtggtgtaccctgtggatgatcatcactttaaggtgatcctgcactatggcacactgg<br>taatcgacggggttacgccgaacatgatcgactatttcggacggccgtatgaaggcatcgccgtgttcgacggc<br>aaaaagatcactgtaacagggaccctgtggaacggcaacaaaattatcgacgagcgcctgatcaaccccga<br>cggctccctgctgttccgagtaaccatcaacggagtgaccggctggcggctgtgcgaacgcattctggcgggc<br>ggcaccggcggcagcggcggcaccggcggcagcatggtcttcacactcgaagatttcgttggggactggcg<br>acagacagccggctacaacctggaccaagtccttgaacagggaggtgtgtccagtttgtttcagaatctcggg<br>gtgtccgtaactccgatccaaaggattgtcctgagcggtgaaaatgggctgaagatcgacatccatgtcatcat<br>cccgtatgaa |
| 1489 | pET28 | $^{CP}$pFAST-DISGG-Nanoluc | ATGGGTGACAGATATTGGGTCTTTGTGAAACGGGTGggcggctccggcggctccgg<br>cggctccggcggcGAGCATGTTGCCTTTGGCAGTGAGGACATCGAGAACACTCT<br>GGCCAATATGGACGACGAACAACTGGATAGGTTGGCCTTTGGCGTAATTCA<br>GCTCGATGGTGACGGGAATATCCTGCTGTACAATGCTGCTGAAGGGGACA<br>TCACTGGCAGAGATCCCAAACAGGTGATTGGGAAGAACTTCTTCAAGGATG<br>TTGCACCTGGAACGGATACTCCCGAGTTTTACGGCAAATTCAAGGAAGGCG<br>CAGCGTCAGGGAATCTGAACACCATGTTCGAATGGACGATACCGACAAGC |

| | | | |
|---|---|---|---|
| | | | AGGGGACCAACCAAGGTCAAGGTGCACTTGAAGAAAGCCCTTTCCgatattagcggcggcatggtcttcacactcgaagatttcgttggggactggcgacagacagccggctacaacctggaccaagtccttgaacagggaggtgtgtccagtttgtttcagaatctcggggtgtccgtaactccgatccaaaggattgtcctgagcggtgaaaatgggctgaagatcgacatccatgtcatcatcccgtatgaaggtctgagcggcgaccaaatgggccagatcgaaaaaatttttaaggtggtgtaccctgtggatgatcatcactttaaggtgatcctgcactatggcacactggtaatcgacggggttacgccgaacatgatcgactatttcggacggccgtatgaaggcatcgccgtgttcgacggcaaaaagatcactgtaacagggaccctgtggaacggcaacaaaattatcgacgagcgcctgatcaaccccgacggctccctgctgttccgagtaaccatcaacggagtgaccggctggcggctgtgcgaacgcattctggcg |
| 1490 | pET28 | Nanoluc-DISGG-$^{CP}$pFAST | atggtcttcacactcgaagatttcgttggggactggcgacagacagccggctacaacctggaccaagtccttgaacagggaggtgtgtccagtttgtttcagaatctcggggtgtccgtaactccgatccaaaggattgtcctgagcggtgaaaatgggctgaagatcgacatccatgtcatcatcccgtatgaaggtctgagcggcgaccaaatgggccagatcgaaaaaatttttaaggtggtgtaccctgtggatgatcatcactttaaggtgatcctgcactatggcacactggtaatcgacggggttacgccgaacatgatcgactatttcggacggccgtatgaaggcatcgccgtgttcgacggcaaaaagatcactgtaacagggaccctgtggaacggcaacaaaattatcgacgagcgcctgatcaaccccgacggctccctgctgttccgagtaaccatcaacggagtgaccggctggcggctgtgcgaacgcattctggcggatattagcggcggcATGGGTGACAGATATTGGGTCTTTGTGAAACGGGTGggcggctccggcggctccggcggctccggcggcGAGCATGTTGCCTTTGGCAGTGAGGACATCGAGAACACTCTGGCCAATATGGACGACGAACAACTGGATAGGTTGGCCTTTGGCGTAATTCAGCTCGATGGTGACGGGAATATCCTGCTGTACAATGCTGCTGAAGGGGACATCACTGGCAGAGATCCCAAACAGGTGATTGGGAAGAACTTCTTCAAGGATGTTGCACCTGGAACGGATACTCCCGAGTTTTACGGCAAATTCAAGGAAGGCGCAGCGTCAGGGAATCTGAACACCATGTTCGAATGGACGATACCGACAAGCAGGGGACCAACCAAGGTCAAGGTGCACTTGAAGAAAGCCCTTTCC |
| 1491 | pET28 | $^{CP}$NanoLuc-DISGG-$^{CP}$pFAST | ggtctgagcggcgaccaaatgggccagatcgaaaaaatttttaaggtggtgtaccctgtggatgatcatcactttaaggtgatcctgcactatggcacactggtaatcgacggggttacgccgaacatgatcgactatttcggacggccgtatgaaggcatcgccgtgttcgacggcaaaaagatcactgtaacagggaccctgtggaacggcaacaaaattatcgacgagcgcctgatcaaccccgacggctccctgctgttccgagtaaccatcaacggagtgaccggctggcggctgtgcgaacgcattctggcgggcggcaccggcggcagcggcggcaccggcggcagcatggtcttcacactcgaagatttcgttggggactggcgacagacagccggctacaacctggaccaagtccttgaacagggaggtgtgtccagtttgtttcagaatctcggggtgtccgtaactccgatccaaaggattgtcctgagcggtgaaaatgggctgaagatcgacatccatgtcatcatcccgtatgaagatattagcggcggcATGGGTGACAGATATTGGGTCTTTGTGAAACGGGTGggcggctccggcggctccggcggctccggcggcGAGCATGTTGCCTTTGGCAGTGAGGACATCGAGAACACTCTGGCCAATATGGACGACGAACAACTGGATAGGTTGGCCTTTGGCGTAATTCAGCTCGATGGTGACGGGAATATCCTGCTGTACAATGCTGCTGAAGGGGACATCACTGGCAGAGATCCCAAACAGGTGATTGGGAAGAACTTCTTCAAGGATGTTGCACCTGGAACGGATACTCCCGAGTTTTACGGCAAATTCAAGGAAGGCGCAGCGTCAGGGAATCTGAACACCATGTTCGAATGGACGATACCGACAAGCAGGGGACCAACCAAGGTCAAGGTGCACTTGAAGAAAGCCCTTTCC |
| 1492 | pET28 | $^{CP}$pFAST-DISGG-$^{CP}$NanoLuc | ATGGGTGACAGATATTGGGTCTTTGTGAAACGGGTGggcggctccggcggctccggcggctccggcggcGAGCATGTTGCCTTTGGCAGTGAGGACATCGAGAACACTCTGGCCAATATGGACGACGAACAACTGGATAGGTTGGCCTTTGGCGTAATTCAGCTCGATGGTGACGGGAATATCCTGCTGTACAATGCTGCTGAAGGGGACATCACTGGCAGAGATCCCAAACAGGTGATTGGGAAGAACTTCTTCAAGGATGTTGCACCTGGAACGGATACTCCCGAGTTTTACGGCAAATTCAAGGAAGGCGCAGCGTCAGGGAATCTGAACACCATGTTCGAATGGACGATACCGACAAGCAGGGGACCAACCAAGGTCAAGGTGCACTTGAAGAAAGCCCTTTCCgatattagcggcggcggtctgagcggcgaccaaatgggccagatcgaaaaaatttttaaggtggtgtaccctgtggatgatcatcactttaaggtgatcctgcactatggcacactggtaatcgacggggttacgccgaacatgatcgactatttcggacggccgtatgaaggcatcgccgtgttcgacggcaaaaagatcactgtaacagggaccctgtggaacggcaacaaaattatcgacgagcgcctgatcaaccccgacggctccctgctgttccgagtaaccatcaacggagtgaccggctggcggctgtgcgaacgcattctggcgggcggcaccggcggcagcggcggcaccggcggcagcatggtcttcacactcgaagatttcgttggggactggcgacagacagccggctacaacctggaccaagtccttgaacagggaggtgtgtccagtttgtttcagaatctcggggtgtccgtaactccgatccaaaggattgtcctgagcggtgaaaatgggctgaagatcgacatccatgtcatcatcccgtatgaa |
| 1493 | pET28 | $^{CP}$NanoLuc | ggtctgagcggcgaccaaatgggccagatcgaaaaaatttttaaggtggtgtaccctgtggatgatcatcactttaaggtgatcctgcactatggcacactggtaatcgacggggttacgccgaacatgatcgactatttcggacggccgtatgaaggcatcgccgtgttcgacggcaaaaagatcactgtaacagggaccctgtggaacggcaacaaaattatcgacgagcgcctgatcaaccccgacggctccctgctgttccgagtaaccatcaacggagtgaccggctggcggctgtgcgaacgcattctggcgggcggcaccggcggcagcggcggcaccggcggcagcatggtcttcacactcgaagatttcgttggggactggcgacagacagccggctacaacctggaccaagtccttgaacagggaggtgtgtccagtttgtttcagaatctcggggtgtccgtaactccgatccaaaggattgtcctgagcggtgaaaatgggctgaagatcgacatccatgtcatcatcccgtatgaa |
| 1546 | pET28 | pFAST-NanoLuc | ATGGAGCATGTTGCCTTTGGCAGTGAGGACATCGAGAACACTCTGGCCAATATGGACGACGAACAACTGGATAGGTTGGCCTTTGGCGTAATTCAGCTCGATGGTGACGGGAATATCCTGCTGTACAATGCTGCTGAAGGGGACATCACTGGCAGAGATCCCAAACAGGTGATTGGGAAGAACTTCTTCAAGGATGTTGCACCTGGAACGGATACTCCCGAGTTTTACGGCAAATTCAAGGAAGGCGCAGCGTCAGGGAATCTGAACACCATGTTCGAATGGACGATACCGACAAGCAGGGGACCAACCAAGGTCAAGGTGCACTTGAAGAAAGCCCTTTCCGGTGACAGATATTGGGTCTTTGTGAAACGGGTGgtcttcacactcgaagatttcgttggggactggcgacagacagccggctacaacctggaccaagtccttgaacagggaggtgtgtccagtttgtttcagaatctcggggtgtccgtaactccgatccaaaggattgtcctgagcggtgaaaatgggctgaagatcgacatccatgtcatcatcccgtatgaa |

| | | | |
|---|---|---|---|
| | | | ggtctgagcggcgaccaaatgggccagatcgaaaaaatttttaaggtggtgtaccctgtggatgatcatcactttaaggtgatcctgcactatggcacactggtaatcgacggggttacgccgaacatgatcgactatttcggacggccgtatgaaggcatcgccgtgttcgacggcaaaaagatcactgtaacagggaccctgtggaacggcaacaaaattatcgacgagcgcctgatcaaccccgacggctccctgctgttccgagtaaccatcaacggagtgaccggctggcggctgtgcgaacgcattctggcg |
| 1547 | pET28 | pFAST-DIS-NanoLuc | ATGGAGCATGTTGCCTTTGGCAGTGAGGACATCGAGAACACTCTGGCCAATATGGACGACGAACAACTGGATAGGTTGGCCTTTGGCGTAATTCAGCTCGATGGTGACGGGAATATCCTGCTGTACAATGCTGCTGAAGGGGACATCACTGGCAGAGATCCCAAACAGGTGATTGGGAAGAACTTCTTCAAGGATGTTGCACCTGGAACGGATACTCCCGAGTTTTACGGCAAATTCAAGGAAGGCGCAGCGTCAGGGAATCTGAACACCATGTTCGAATGGACGATACCGACAAGCAGGGGACCAACCAAGGTCAAGGTGCACTTGAAGAAAGCCCTTTCCGGTGACAGATATTGGGTCTTTGTGAAACGGGTGgatattagcgtcttcacactcgaagatttcgttggggactggcgacagacagccggctacaacctggaccaagtccttgaacagggaggtgtgtccagtttgtttcagaatctcggggtgtccgtaactccgatccaaaggattgtcctgagcggtgaaaatgggctgaagatcgacatccatgtcatcatcccgtatgaaggtctgagcggcgaccaaatgggccagatcgaaaaaatttttaaggtggtgtaccctgtggatgatcatcactttaaggtgatcctgcactatggcacactggtaatcgacggggttacgccgaacatgatcgactatttcggacggccgtatgaaggcatcgccgtgttcgacggcaaaaagatcactgtaacagggaccctgtggaacggcaacaaaattatcgacgagcgcctgatcaaccccgacggctccctgctgttccgagtaaccatcaacggagtgaccggctggcggctgtgcgaacgcattctggcg |
| 1548 | pET28 | pFAST-DISGGDIS-NanoLuciferase | ATGGAGCATGTTGCCTTTGGCAGTGAGGACATCGAGAACACTCTGGCCAATATGGACGACGAACAACTGGATAGGTTGGCCTTTGGCGTAATTCAGCTCGATGGTGACGGGAATATCCTGCTGTACAATGCTGCTGAAGGGGACATCACTGGCAGAGATCCCAAACAGGTGATTGGGAAGAACTTCTTCAAGGATGTTGCACCTGGAACGGATACTCCCGAGTTTTACGGCAAATTCAAGGAAGGCGCAGCGTCAGGGAATCTGAACACCATGTTCGAATGGACGATACCGACAAGCAGGGGACCAACCAAGGTCAAGGTGCACTTGAAGAAAGCCCTTTCCGGTGACAGATATTGGGTCTTTGTGAAACGGGTGgatattagcggcggcgatattagcgtcttcacactcgaagatttcgttggggactggcgacagacagccggctacaacctggaccaagtccttgaacagggaggtgtgtccagtttgtttcagaatctcggggtgtccgtaactccgatccaaaggattgtcctgagcggtgaaaatgggctgaagatcgacatccatgtcatcatcccgtatgaaggtctgagcggcgaccaaatgggccagatcgaaaaaatttttaaggtggtgtaccctgtggatgatcatcactttaaggtgatcctgcactatggcacactggtaatcgacggggttacgccgaacatgatcgactatttcggacggccgtatgaaggcatcgccgtgttcgacggcaaaaagatcactgtaacagggaccctgtggaacggcaacaaaattatcgacgagcgcctgatcaaccccgacggctccctgctgttccgagtaaccatcaacggagtgaccggctggcggctgtgcgaacgcattctggcg |
| 1549 | pET28 | pFAST-DISGGDISGGD-NanoLuciferase | ATGGAGCATGTTGCCTTTGGCAGTGAGGACATCGAGAACACTCTGGCCAATATGGACGACGAACAACTGGATAGGTTGGCCTTTGGCGTAATTCAGCTCGATGGTGACGGGAATATCCTGCTGTACAATGCTGCTGAAGGGGACATCACTGGCAGAGATCCCAAACAGGTGATTGGGAAGAACTTCTTCAAGGATGTTGCACCTGGAACGGATACTCCCGAGTTTTACGGCAAATTCAAGGAAGGCGCAGCGTCAGGGAATCTGAACACCATGTTCGAATGGACGATACCGACAAGCAGGGGACCAACCAAGGTCAAGGTGCACTTGAAGAAAGCCCTTTCCGGTGACAGATATTGGGTCTTTGTGAAACGGGTGgatattagcggcggcgatattagcggcggcgatgtcttcacactcgaagatttcgttggggactggcgacagacagccggctacaacctggaccaagtccttgaacagggaggtgtgtccagtttgtttcagaatctcggggtgtccgtaactccgatccaaaggattgtcctgagcggtgaaaatgggctgaagatcgacatccatgtcatcatcccgtatgaaggtctgagcggcgaccaaatgggccagatcgaaaaaatttttaaggtggtgtaccctgtggatgatcatcactttaaggtgatcctgcactatggcacactggtaatcgacggggttacgccgaacatgatcgactatttcggacggccgtatgaaggcatcgccgtgttcgacggcaaaaagatcactgtaacagggaccctgtggaacggcaacaaaattatcgacgagcgcctgatcaaccccgacggctccctgctgttccgagtaaccatcaacggagtgaccggctggcggctgtgcgaacgcattctggcg |
| 1566 | pET28 | $^{C}$FAST-$^{CP}$NanoLuc-$^{N}$FAST | atgGGTGACAGATATTGGGTCTTTGTGAAACGGGTGggtctgagcggcgaccaaatgggccagatcgaaaaaatttttaaggtggtgtaccctgtggatgatcatcactttaaggtgatcctgcactatggcacactggtaatcgacggggttacgccgaacatgatcgactatttcggacggccgtatgaaggcatcgccgtgttcgacggcaaaaagatcactgtaacagggaccctgtggaacggcaacaaaattatcgacgagcgcctgatcaaccccgacggctccctgctgttccgagtaaccatcaacggagtgaccggctggcggctgtgcgaacgcattctggcgggcggcaccggcggcagcggcggcaccggcggcagcatggtcttcacactcgaagatttcgttggggactggcgacagacagccggctacaacctggaccaagtccttgaacagggaggtgtgtccagtttgtttcagaatctcggggtgtccgtaactccgatccaaaggattgtcctgagcggtgaaaatgggctgaagatcgacatccatgtcatcatcccgtatgaaGAGCATGTTGCCTTTGGCAGTGAGGACATCGAGAACACTCTGGCCAATATGGACGACGAACAACTGGATAGGTTGGCCTTTGGCGTAATTCAGCTCGATGGTGACGGGAATATCCTGCTGTACAATGCTGCTGAAGGGGACATCACTGGCAGAGATCCCAAACAGGTGATTGGGAAGAACTTCTTCAAGGATGTTGCACCTGGAACGGATACTCCCGAGTTTTACGGCAAATTCAAGGAAGGCGCAGCGTCAGGGAATCTGAACACCATGTTCGAATGGACGATACCGACAAGCAGGGGACCAACCAAGGTCAAGGTGCACTTGAAGAAAGCCCTTTCC |
| 1567 | pET28 | $^{C}$FAST-NanoLuc-$^{N}$FAST | atgGGTGACAGATATTGGGTCTTTGTGAAACGGGTGatggtcttcacactcgaagatttcgttggggactggcgacagacagccggctacaacctggaccaagtccttgaacagggaggtgtgtccagtttgtttcagaatctcggggtgtccgtaactccgatccaaaggattgtcctgagcggtgaaaatgggctgaagatcgacatccatgtcatcatcccgtatgaaggtctgagcggcgaccaaatgggccagatcgaaaaaatttttaaggtggtgtaccctgtggatgatcatcactttaaggtgatcctgcactatggcacactggtaatcgacggggttacgccgaacatgatcgactatttcggacggccgtatgaaggcatcgccgtgttcgacggcaaaaagatcactgtaacagggaccctgtggaacggcaacaaaattatcgacgagcgcctgatcaaccccgacggctccctgctgttccgagtaaccatcaacggagtgaccggctggcggctgtgcgaacgcattctggcgGAGCATGTTGCCTTTGGCAGTGAGGACATCGAGAACACTCTGGCCAATATGGACGACGAACAACTGGATAGGTTGGCCTTTGGCGTAATTCAGCTCGATGGTGACGGGAATATCCTGC |

| | | | |
|---|---|---|---|
| | | | TGTACAATGCTGCTGAAGGGGACATCACTGGCAGAGATCCCAAACAGGTGATTGGGAAGAACTTCTTCAAGGATGTTGCACCTGGAACGGATACTCCCGAGTTTTACGGCAAATTCAAGGAAGGCGCAGCGTCAGGGAATCTGAACACCATGTTCGAATGGACGATACCGACAAGCAGGGGACCAACCAAGGTCAAGGTGCACTTGAAGAAAGCCCTTTCC |
| 1568 | pET28 | $^{C}$Nanoluc-$^{CP}$pFAST-$^{N}$NanoLuc | ggtctgagcggcgaccaaatgggccagatcgaaaaaatttttaaggtggtgtaccctgtggatgatcatcactttaaggtgatcctgcactatggcacactggtaatcgacggggttacgccgaacatgatcgactatttcggacggccgtatgaaggcatcgccgtgttcgacggcaaaaagatcactgtaacagggaccctgtggaacggcaacaaaattatcgacgagcgcctgatcaaccccgacggctccctgctgttccgagtaaccatcaacggagtgaccggctggcggctgtgcgaacgcattctggcgATGGGTGACAGATATTGGGTCTTTGTGAAACGGGTGggcggctccggcggctccggcggctccggcggcGAGCATGTTGCCTTTGGCAGTGAGGACATCGAGAACACTCTGGCCAATATGGACGACGAACAACTGGATAGGTTGGCCTTTGGCGTAATTCAGCTCGATGGTGACGGGAATATCCTGCTGTACAATGCTGCTGAAGGGGACATCACTGGCAGAGATCCCAAACAGGTGATTGGGAAGAACTTCTTCAAGGATGTTGCACCTGGAACGGATACTCCCGAGTTTTACGGCAAATTCAAGGAAGGCGCAGCGTCAGGGAATCTGAACACCATGTTCGAATGGACGATACCGACAAGCAGGGGACCAACCAAGGTCAAGGTGCACTTGAAGAAAGCCCTTTCCatggtcttcacactcgaagatttcgttggggactggcgacagacagccggctacaacctggaccaagtccttgaacagggaggtgtgtccagtttgtttcagaatctcggggtgtccgtaactccgatccaaaggattgtcctgagcggtgaaaatgggctgaagatcgacatccatgtcatcatcccgtatgaa |
| 1569 | pET28 | $^{C}$Nanoluc-pFAST-$^{N}$NanoLuc | ggtctgagcggcgaccaaatgggccagatcgaaaaaatttttaaggtggtgtaccctgtggatgatcatcactttaaggtgatcctgcactatggcacactggtaatcgacggggttacgccgaacatgatcgactatttcggacggccgtatgaaggcatcgccgtgttcgacggcaaaaagatcactgtaacagggaccctgtggaacggcaacaaaattatcgacgagcgcctgatcaaccccgacggctccctgctgttccgagtaaccatcaacggagtgaccggctggcggctgtgcgaacgcattctggcgGAGCATGTTGCCTTTGGCAGTGAGGACATCGAGAACACTCTGGCCAATATGGACGACGAACAACTGGATAGGTTGGCCTTTGGCGTAATTCAGCTCGATGGTGACGGGAATATCCTGCTGTACAATGCTGCTGAAGGGGACATCACTGGCAGAGATCCCAAACAGGTGATTGGGAAGAACTTCTTCAAGGATGTTGCACCTGGAACGGATACTCCCGAGTTTTACGGCAAATTCAAGGAAGGCGCAGCGTCAGGGAATCTGAACACCATGTTCGAATGGACGATACCGACAAGCAGGGGACCAACCAAGGTCAAGGTGCACTTGAAGAAAGCCCTTTCCGGTGACAGATATTGGGTCTTTGTGAAACGGGTGatggtcttcacactcgaagatttcgttggggactggcgacagacagccggctacaacctggaccaagtccttgaacagggaggtgtgtccagtttgtttcagaatctcggggtgtccgtaactccgatccaaaggattgtcctgagcggtgaaaatgggctgaagatcgacatccatgtcatcatcccgtatgaa |
| 1570 | pET28 | $^{N}$Nanoluc-$^{CP}$pFAST-$^{C}$NanoLuc | atggtcttcacactcgaagatttcgttggggactggcgacagacagccggctacaacctggaccaagtccttgaacagggaggtgtgtccagtttgtttcagaatctcggggtgtccgtaactccgatccaaaggattgtcctgagcggtgaaaatgggctgaagatcgacatccatgtcatcatcccgtatgaaATGGGTGACAGATATTGGGTCTTTGTGAAACGGGTGggcggctccggcggctccggcggctccggcggcGAGCATGTTGCCTTTGGCAGTGAGGACATCGAGAACACTCTGGCCAATATGGACGACGAACAACTGGATAGGTTGGCCTTTGGCGTAATTCAGCTCGATGGTGACGGGAATATCCTGCTGTACAATGCTGCTGAAGGGGACATCACTGGCAGAGATCCCAAACAGGTGATTGGGAAGAACTTCTTCAAGGATGTTGCACCTGGAACGGATACTCCCGAGTTTTACGGCAAATTCAAGGAAGGCGCAGCGTCAGGGAATCTGAACACCATGTTCGAATGGACGATACCGACAAGCAGGGGACCAACCAAGGTCAAGGTGCACTTGAAGAAAGCCCTTTCCggtctgagcggcgaccaaatgggccagatcgaaaaaatttttaaggtggtgtaccctgtggatgatcatcactttaaggtgatcctgcactatggcacactggtaatcgacggggttacgccgaacatgatcgactatttcggacggccgtatgaaggcatcgccgtgttcgacggcaaaaagatcactgtaacagggaccctgtggaacggcaacaaaattatcgacgagcgcctgatcaaccccgacggctccctgctgttccgagtaaccatcaacggagtgaccggctggcggctgtgcgaacgcattctggcg |
| 1571 | pET28 | $^{N}$Nanoluc-pFAST-$^{C}$NanoLuc | atggtcttcacactcgaagatttcgttggggactggcgacagacagccggctacaacctggaccaagtccttgaacagggaggtgtgtccagtttgtttcagaatctcggggtgtccgtaactccgatccaaaggattgtcctgagcggtgaaaatgggctgaagatcgacatccatgtcatcatcccgtatgaaATGGAGCATGTTGCCTTTGGCAGTGAGGACATCGAGAACACTCTGGCCAATATGGACGACGAACAACTGGATAGGTTGGCCTTTGGCGTAATTCAGCTCGATGGTGACGGGAATATCCTGCTGTACAATGCTGCTGAAGGGGACATCACTGGCAGAGATCCCAAACAGGTGATTGGGAAGAACTTCTTCAAGGATGTTGCACCTGGAACGGATACTCCCGAGTTTTACGGCAAATTCAAGGAAGGCGCAGCGTCAGGGAATCTGAACACCATGTTCGAATGGACGATACCGACAAGCAGGGGACCAACCAAGGTCAAGGTGCACTTGAAGAAAGCCCTTTCCGGTGACAGATATTGGGTCTTTGTGAAACGGGTGggtctgagcggcgaccaaatgggccagatcgaaaaaatttttaaggtggtgtaccctgtggatgatcatcactttaaggtgatcctgcactatggcacactggtaatcgacggggttacgccgaacatgatcgactatttcggacggccgtatgaaggcatcgccgtgttcgacggcaaaaagatcactgtaacagggaccctgtggaacggcaacaaaattatcgacgagcgcctgatcaaccccgacggctccctgctgttccgagtaaccatcaacggagtgaccggctggcggctgtgcgaacgcattctggcg |
| 1572 | pET28 | $^{N}$pFAST-$^{CP}$NanoLuc-$^{C}$pFAST | atgGAGCATGTTGCCTTTGGCAGTGAGGACATCGAGAACACTCTGGCCAATATGGACGACGAACAACTGGATAGGTTGGCCTTTGGCGTAATTCAGCTCGATGGTGACGGGAATATCCTGCTGTACAATGCTGCTGAAGGGGACATCACTGGCAGAGATCCCAAACAGGTGATTGGGAAGAACTTCTTCAAGGATGTTGCACCTGGAACGGATACTCCCGAGTTTTACGGCAAATTCAAGGAAGGCGCAGCGTCAGGGAATCTGAACACCATGTTCGAATGGACGATACCGACAAGCAGGGGACCAACCAAGGTCAAGGTGCACTTGAAGAAAGCCCTTTCCggtctgagcggcgaccaaatgggccagatcgaaaaaatttttaaggtggtgtaccctgtggatgatcatcactttaaggtgatcctgcactatggcacactggtaatcgacggggttacgccgaacatgatcgactatttcggacggccgtatgaaggcatcgccgtgttcgacggcaaaaagatcactgtaacagggaccctgtggaacggcaacaaaattatcgacgagcgcctga |

| | | | |
|---|---|---|---|
| | | | tcaaccccgacggctccctgctgttccgagtaaccatcaacggagtgaccggctggcggctgtgcgaacgcat<br>tctggcgggcggcaccggcggcagcggcggcaccggcggcagcatggtcttcacactcgaagatttcgttgg<br>ggactggcgacagacagccggctacaacctggaccaagtccttgaacagggaggtgtgtccagtttgtttcag<br>aatctcggggtgtccgtaactccgatccaaaggattgtcctgagcggtgaaaatgggctgaagatcgacatcc<br>atgtcatcatcccgtatgaaGGTGACAGATATTGGGTCTTTGTGAAACGGGTG |
| 1573 | pET28 | $^{N}$pFAST-<br>NanoLuc-<br>$^{C}$pFAST | atgGAGCATGTTGCCTTTGGCAGTGAGGACATCGAGAACACTCTGGCCAATA<br>TGGACGACGAACAACTGGATAGGTTGGCCTTTGGCGTAATTCAGCTCGATG<br>GTGACGGGAATATCCTGCTGTACAATGCTGCTGAAGGGGACATCACTGGC<br>AGAGATCCCAAACAGGTGATTGGGAAGAACTTCTTCAAGGATGTTGCACCT<br>GGAACGGATACTCCCGAGTTTTACGGCAAATTCAAGGAAGGCGCAGCGTC<br>AGGGAATCTGAACACCATGTTCGAATGGACGATACCGACAAGCAGGGGAC<br>CAACCAAGGTCAAGGTGCACTTGAAGAAAGCCCTTTCCatggtcttcacactcgaag<br>atttcgttggggactggcgacagacagccggctacaacctggaccaagtccttgaacagggaggtgtgtcca<br>gtttgtttcagaatctcggggtgtccgtaactccgatccaaaggattgtcctgagcggtgaaaatgggctgaagat<br>cgacatccatgtcatcatcccgtatgaaggtctgagcggcgaccaaatgggccagatcgaaaaaatttttaag<br>gtggtgtaccctgtggatgatcatcactttaaggtgatcctgcactatggcacactggtaatcgacggggttacgc<br>cgaacatgatcgactatttcggacggccgtatgaaggcatcgccgtgttcgacggcaaaaagatcactgtaac<br>agggaccctgtggaacggcaacaaaattatcgacgagcgcctgatcaaccccgacggctccctgctgttccg<br>agtaaccatcaacggagtgaccggctggcggctgtgcgaacgcattctggcgGGTGACAGATATTG<br>GGTCTTTGTGAAACGGGTG |
| 1621 | pET28 | pFAST-DISGGD-<br>NanoLuc | ATGGAGCATGTTGCCTTTGGCAGTGAGGACATCGAGAACACTCTGGCCAAT<br>ATGGACGACGAACAACTGGATAGGTTGGCCTTTGGCGTAATTCAGCTCGAT<br>GGTGACGGGAATATCCTGCTGTACAATGCTGCTGAAGGGGACATCACTGG<br>CAGAGATCCCAAACAGGTGATTGGGAAGAACTTCTTCAAGGATGTTGCACC<br>TGGAACGGATACTCCCGAGTTTTACGGCAAATTCAAGGAAGGCGCAGCGT<br>CAGGGAATCTGAACACCATGTTCGAATGGACGATACCGACAAGCAGGGGA<br>CCAACCAAGGTCAAGGTGCACTTGAAGAAAGCCCTTTCCGGTGACAGATAT<br>TGGGTCTTTGTGAAACGGGTGgatattagcggcggcgatgtcttcacactcgaagatttcgttggg<br>gactggcgacagacagccggctacaacctggaccaagtccttgaacagggaggtgtgtccagtttgtttcaga<br>atctcggggtgtccgtaactccgatccaaaggattgtcctgagcggtgaaaatgggctgaagatcgacatccat<br>gtcatcatcccgtatgaaggtctgagcggcgaccaaatgggccagatcgaaaaaatttttaaggtggtgtaccc<br>tgtggatgatcatcactttaaggtgatcctgcactatggcacactggtaatcgacggggttacgccgaacatgat<br>cgactatttcggacggccgtatgaaggcatcgccgtgttcgacggcaaaaagatcactgtaacagggaccctg<br>tggaacggcaacaaaattatcgacgagcgcctgatcaaccccgacggctccctgctgttccgagtaaccatca<br>acggagtgaccggctggcggctgtgcgaacgcattctggcg |
| 1622 | pET28 | pFAST-<br>DISGGDI-<br>NanoLuc | ATGGAGCATGTTGCCTTTGGCAGTGAGGACATCGAGAACACTCTGGCCAAT<br>ATGGACGACGAACAACTGGATAGGTTGGCCTTTGGCGTAATTCAGCTCGAT<br>GGTGACGGGAATATCCTGCTGTACAATGCTGCTGAAGGGGACATCACTGG<br>CAGAGATCCCAAACAGGTGATTGGGAAGAACTTCTTCAAGGATGTTGCACC<br>TGGAACGGATACTCCCGAGTTTTACGGCAAATTCAAGGAAGGCGCAGCGT<br>CAGGGAATCTGAACACCATGTTCGAATGGACGATACCGACAAGCAGGGGA<br>CCAACCAAGGTCAAGGTGCACTTGAAGAAAGCCCTTTCCGGTGACAGATAT<br>TGGGTCTTTGTGAAACGGGTGgatattagcggcggcgatattgtcttcacactcgaagatttcgttg<br>gggactggcgacagacagccggctacaacctggaccaagtccttgaacagggaggtgtgtccagtttgtttca<br>gaatctcggggtgtccgtaactccgatccaaaggattgtcctgagcggtgaaaatgggctgaagatcgacatc<br>catgtcatcatcccgtatgaaggtctgagcggcgaccaaatgggccagatcgaaaaaatttttaaggtggtgta<br>ccctgtggatgatcatcactttaaggtgatcctgcactatggcacactggtaatcgacggggttacgccgaacat<br>gatcgactatttcggacggccgtatgaaggcatcgccgtgttcgacggcaaaaagatcactgtaacagggac<br>cctgtggaacggcaacaaaattatcgacgagcgcctgatcaaccccgacggctccctgctgttccgagtaacc<br>atcaacggagtgaccggctggcggctgtgcgaacgcattctggcg |
| 1623 | pET28 | pFAST-<br>DISGGDISG-<br>NanoLuc | ATGGAGCATGTTGCCTTTGGCAGTGAGGACATCGAGAACACTCTGGCCAAT<br>ATGGACGACGAACAACTGGATAGGTTGGCCTTTGGCGTAATTCAGCTCGAT<br>GGTGACGGGAATATCCTGCTGTACAATGCTGCTGAAGGGGACATCACTGG<br>CAGAGATCCCAAACAGGTGATTGGGAAGAACTTCTTCAAGGATGTTGCACC<br>TGGAACGGATACTCCCGAGTTTTACGGCAAATTCAAGGAAGGCGCAGCGT<br>CAGGGAATCTGAACACCATGTTCGAATGGACGATACCGACAAGCAGGGGA<br>CCAACCAAGGTCAAGGTGCACTTGAAGAAAGCCCTTTCCGGTGACAGATAT<br>TGGGTCTTTGTGAAACGGGTGgatattagcggcggcgatattagcggcgtcttcacactcgaag<br>atttcgttggggactggcgacagacagccggctacaacctggaccaagtccttgaacagggaggtgtgtcca<br>gtttgtttcagaatctcggggtgtccgtaactccgatccaaaggattgtcctgagcggtgaaaatgggctgaagat<br>cgacatccatgtcatcatcccgtatgaaggtctgagcggcgaccaaatgggccagatcgaaaaaatttttaag<br>gtggtgtaccctgtggatgatcatcactttaaggtgatcctgcactatggcacactggtaatcgacggggttacgc<br>cgaacatgatcgactatttcggacggccgtatgaaggcatcgccgtgttcgacggcaaaaagatcactgtaac<br>agggaccctgtggaacggcaacaaaattatcgacgagcgcctgatcaaccccgacggctccctgctgttccg<br>agtaaccatcaacggagtgaccggctggcggctgtgcgaacgcattctggcg |
| 1624 | pET28 | pFAST-<br>DISGGDISGG-<br>NanoLuc | ATGGAGCATGTTGCCTTTGGCAGTGAGGACATCGAGAACACTCTGGCCAAT<br>ATGGACGACGAACAACTGGATAGGTTGGCCTTTGGCGTAATTCAGCTCGAT<br>GGTGACGGGAATATCCTGCTGTACAATGCTGCTGAAGGGGACATCACTGG<br>CAGAGATCCCAAACAGGTGATTGGGAAGAACTTCTTCAAGGATGTTGCACC<br>TGGAACGGATACTCCCGAGTTTTACGGCAAATTCAAGGAAGGCGCAGCGT<br>CAGGGAATCTGAACACCATGTTCGAATGGACGATACCGACAAGCAGGGGA<br>CCAACCAAGGTCAAGGTGCACTTGAAGAAAGCCCTTTCCGGTGACAGATAT<br>TGGGTCTTTGTGAAACGGGTGgatattagcggcggcgatattagcggcggcgtcttcacactcg<br>aagatttcgttggggactggcgacagacagccggctacaacctggaccaagtccttgaacagggaggtgtgt<br>ccagtttgtttcagaatctcggggtgtccgtaactccgatccaaaggattgtcctgagcggtgaaaatgggctga |

| | | | |
|---|---|---|---|
| | | | agatcgacatccatgtcatcatcccgtatgaaggtctgagcggcgaccaaatgggccagatcgaaaaaatttttaaggtggtgtaccctgtggatgatcatcactttaaggtgatcctgcactatggcacactggtaatcgacggggttacgccgaacatgatcgactatttcggacggccgtatgaaggcatcgccgtgttcgacggcaaaaagatcactgtaacagggaccctgtggaacggcaacaaaattatcgacgagcgcctgatcaaccccgacggctccctgctgttccgagtaaccatcaacggagtgaccggctggcggctgtgcgaacgcattctggcg |
| 1654 | CMV | LumiFAST | ATGGAGCATGTTGCCTTTGGCAGTGAGGACATCGAGAACACTCTGGCCAATATGGACGACGAACAACTGGATAGGTTGGCCTTTGGCGTAATTCAGCTCGATGGTGACGGGAATATCCTGCTGTACAATGCTGCTGAAGGGGACATCACTGGCAGAGATCCCAAACAGGTGATTGGGAAGAACTTCTTCAAGGATGTTGCACCTGGAACGGATACTCCCGAGTTTTACGGCAAATTCAAGGAAGGCGCAGCGTCAGGGAATCTGAACACCATGTTCGAATGGACGATACCGACAAGCAGGGGACCAACCAAGGTCAAGGTGCACTTGAAGAAAGCCCTTTCCGGTGACAGATATTGGGTCTTTGTGAAACGGGTGgatattagcggcggcgatattgtcttcacactcgaagatttcgttggggactggcgacagacagccggctacaacctggaccaagtccttgaacagggaggtgtgtccagtttgtttcagaatctcggggtgtccgtaactccgatccaaaggattgtcctgagcggtgaaaatgggctgaagatcgacatccatgtcatcatcccgtatgaaggtctgagcggcgaccaaatgggccagatcgaaaaaatttttaaggtggtgtaccctgtggatgatcatcactttaaggtgatcctgcactatggcacactggtaatcgacggggttacgccgaacatgatcgactatttcggacggccgtatgaaggcatcgccgtgttcgacggcaaaaagatcactgtaacagggaccctgtggaacggcaacaaaattatcgacgagcgcctgatcaaccccgacggctccctgctgttccgagtaaccatcaacggagtgaccggctggcggctgtgcgaacgcattctggcg |
| 1790 | CMV | H2B-LumiFAST | ATGCCCGAACCTGCGAAGTCAGCGCCCGCTCCCAAAAAAGGCTCTAAAAAAGCTGTCGCCAAGACCCAGAAGAAGGGGGATAAGAAAAGGCGTAAGACCAGGAAAGAGAGTTACGCCATTTACGTGTACAAAGTACTAAAACAAGTCCACCCGGACACTGGCATCTCCTCAAAGGCGATGGGCATTATGAACTCATTTGTAAACGACATCTTCGAGCGCATCGCCGGAGAAGCGTCGCGCCTGGCGCATTACAACAAGCGCTCCACTATCACATCCCGGGAGATCCAGACGGCCGTGCGCCTGCTCCTGCCCGGAGAACTGGCCAAACACGCTGTGTCTGAGGGCACAAAGGCCGTGACCAAGTACACCAGCTCCAAGggcggaggctccggaggcggatctgccaccATGGAGCATGTTGCCTTTGGCAGTGAGGACATCGAGAACACTCTGGCCAATATGGACGACGAACAACTGGATAGGTTGGCCTTTGGCGTAATTCAGCTCGATGGTGACGGGAATATCCTGCTGTACAATGCTGCTGAAGGGGACATCACTGGCAGAGATCCCAAACAGGTGATTGGGAAGAACTTCTTCAAGGATGTTGCACCTGGAACGGATACTCCCGAGTTTTACGGCAAATTCAAGGAAGGCGCAGCGTCAGGGAATCTGAACACCATGTTCGAATGGACGATACCGACAAGCAGGGGACCAACCAAGGTCAAGGTGCACTTGAAGAAAGCCCTTTCCGGTGACAGATATTGGGTCTTTGTGAAACGGGTGgatattagcggcggcgatattgtcttcacactcgaagatttcgttggggactggcgacagacagccggctacaacctggaccaagtccttgaacagggaggtgtgtccagtttgtttcagaatctcggggtgtccgtaactccgatccaaaggattgtcctgagcggtgaaaatgggctgaagatcgacatccatgtcatcatcccgtatgaaggtctgagcggcgaccaaatgggccagatcgaaaaaatttttaaggtggtgtaccctgtggatgatcatcactttaaggtgatcctgcactatggcacactggtaatcgacggggttacgccgaacatgatcgactatttcggacggccgtatgaaggcatcgccgtgttcgacggcaaaaagatcactgtaacagggaccctgtggaacggcaacaaaattatcgacgagcgcctgatcaaccccgacggctccctgctgttccgagtaaccatcaacggagtgaccggctggcggctgtgcgaacgcattctggcg |
| 1798 | CMV | H2B-NanoLuc-iRFP670 | ATGCCCGAACCTGCGAAGTCAGCGCCCGCTCCCAAAAAAGGCTCTAAAAAAGCTGTCGCCAAGACCCAGAAGAAGGGGGATAAGAAAAGGCGTAAGACCAGGAAAGAGAGTTACGCCATTTACGTGTACAAAGTACTAAAACAAGTCCACCCGGACACTGGCATCTCCTCAAAGGCGATGGGCATTATGAACTCATTTGTAAACGACATCTTCGAGCGCATCGCCGGAGAAGCGTCGCGCCTGGCGCATTACAACAAGCGCTCCACTATCACATCCCGGGAGATCCAGACGGCCGTGCGCCTGCTCCTGCCCGGAGAACTGGCCAAACACGCTGTGTCTGAGGGCACAAAGGCCGTGACCAAGTACACCAGCTCCAAGggcggaggctccggaggcggatctgccaccATGGTCTTCACACTCGAAGATTTCGTTGGGGACTGGCGACAGACAGCCGGCTACAACCTGGACCAAGTCCTTGAACAGGGAGGTGTGTCCAGTTTGTTTCAGAATCTCGGGGTGTCCGTAACTCCGATCCAAAGGATTGTCCTGAGCGGTGAAAATGGGCTGAAGATCGACATCCATGTCATCATCCCGTATGAAGGTCTGAGCGGCGACCAAATGGGCCAGATCGAAAAAATTTTTAAGGTGGTGTACCCTGTGGATGATCATCACTTTAAGGTGATCCTGCACTATGGCACACTGGTAATCGACGGGGTTACGCCGAACATGATCGACTATTTCGGACGGCCGTATGAAGGCATCGCCGTGTTCGACGGCAAAAAGATCACTGTAACAGGGACCCTGTGGAACGGCAACAAAATTATCGACGAGCGCCTGATCAACCCCGACGGCTCCCTGCTGTTCCGAGTAACCATCAACGGAGTGACCGGCTGGCGGCTGTGCGAACGCATTCTGGCGTAACTCGAGGACTACAAGGACGACGACGACAAGCCCGGGatccgcccctctccctcccccccccctaacgttactggccgaagccgcttggaataaggccggtgtgcgtttgtctatatgttattttccaccatattgccgtcttttggcaatgtgagggcccggaaacctggccctgtcttcttgacgagcattcctaggggtctttcccctctcgccaaaggaatgcaaggtctgttgaatgtcgtgaaggaagcagttcctctggaagcttcttgaagacaaacaacgtctgtagcgaccctttgcaggcagcggaaccccccacctggcgacaggtgcctctgcggccaaaagccacgtgtataagatacacctgcaaaggcggcacaaccccagtgccacgttgtgagttggatagttgtggaaagagtcaaatggctctcctcaagcgtattcaacaaggggctgaaggatgcccagaaggtaccccattgtatgggatctgatctggggcctcggtgcacatgctttacatgtgtttagtcgaggttaaaaaaacgtctaggccccccgaaccacggggacgtggttttcctttgaaaaacacgatgataatatggCCACaaccATGcgatcgTACCCATACGATGTTCCAGATTACGCTgaattcatggcgcgtaaggtcgatctcacctcctgcgatcgcgagccgatccacatccccggcagcattcagccgtgcggctgcctgctagcctgcgacgcgcaggcggtgcggatcacgcgcattacgggaaatgccggcgcgttctttggacgcgaaactccgcgggtcggtgagctactcgccgattacttcggcgagaccgaagcccatgcgctgcgcaacgcactggcgcagtcttccgatccaaagcgaccggcgctgatcttcggttggcgcgacggcctgaccggccgcaccttcgacatctcactgcatcgccatg |

| | | | |
|---|---|---|---|
| | | | acggtacatcgatcatcgagttcgagcctgcggcggccgaacaggccgacaatccgctgcggctgacgcggcagatcatcgcgcgcaccaaagaactgaagtcgctcgaagagatggccgcacgggtgccgcgctatctgcaggcgatgctcggctatcaccgcgtgatgttgtaccgcttcgcggacgacggctccgggatggtgatcggcgaggcgaagcgcagcgacctcgagagctttctcggtcagcactttccggcgtcgctggtcccgcagcaggcgcggctactgtacttgaagaacgcgatccgcgtggtctcggattcgcgcggcatcagcagccggatcgtgcccgagcacgacgcctccggcgccgcgctcgatctgtcgttcgcgcacctgcgcagcatctcgccctgccatctcgaatttctgcggaacatgggcgtcagcgcctcgatgtcgctgtcgatcatcattgacggcacgctatggggattgatcatctgtcatcattacgagccgcgtgccgtgccgatggcgcagcgcgtcgcggccgaaatgttcgccgacttcttatcgctgcacttcaccgccgcccaccaccaacgc |
| 1799 | CMV | Tom20-NanoLuc-iRFP670 | atggtgggtcggaacagcgccatcgccgcgggcgtgtgcggtgccctcttcatagggtactgcatctactttgaccgcaaaagacgaagtgaccccaacttcagatctgccaccATGGTCTTCACACTCGAAGATTTCGTTGGGGACTGGCGACAGACAGCCGGCTACAACCTGGACCAAGTCCTTGAACAGGGAGGTGTGTCCAGTTTGTTTCAGAATCTCGGGGTGTCCGTAACTCCGATCCAAAGGATTGTCCTGAGCGGTGAAAATGGGCTGAAGATCGACATCCATGTCATCATCCCGTATGAAGGTCTGAGCGGCGACCAAATGGGCCAGATCGAAAAAATTTTTAAGGTGGTGTACCCTGTGGATGATCATCACTTTAAGGTGATCCTGCACTATGGCACACTGGTAATCGACGGGGTTACGCCGAACATGATCGACTATTTCGGACGGCCGTATGAAGGCATCGCCGTGTTCGACGGCAAAAAGATCACTGTAACAGGGACCCTGTGGAACGGCAACAAAATTATCGACGAGCGCCTGATCAACCCCGACGGCTCCCTGCTGTTCCGAGTAACCATCAACGGAGTGACCGGCTGGCGGCTGTGCGAACGCATTCTGGCGTAACTCGAGGACTACAAGGACGACGACGACAAGCCCGGGatccgcccctctccctcccccccccctaacgttactggccgaagccgcttggaataaggccggtgtgcgtttgtctatatgttattttccaccatattgccgtcttttggcaatgtgagggcccggaaacctggccctgtcttcttgacgagcattcctaggggtctttcccctctcgccaaaggaatgcaaggtctgttgaatgtcgtgaaggaagcagttcctctggaagcttcttgaagacaaacaacgtctgtagcgaccctttgcaggcagcggaaccccccacctggcgacaggtgcctctgcggccaaaagccacgtgtataagatacacctgcaaaggcggcacaaccccagtgccacgttgtgagttggatagttgtggaaagagtcaaatggctctcctcaagcgtattcaacaaggggctgaaggatgcccagaaggtaccccattgtatgggatctgatctggggcctcggtgcacatgctttacatgtgtttagtcgaggttaaaaaaacgtctaggccccccgaaccacggggacgtggttttcctttgaaaaacacgatgataatatggCCACaaccATGcgatcgTACCCATACGATGTTCCAGATTACGCTgaattcatggcgcgtaaggtcgatctcacctcctgcgatcgcgagccgatccacatccccggcagcattcagccgtgcggctgcctgctagcctgcgacgcgcaggcggtgcggatcacgcgcattacggaaaatgccggcgcgttctttggacgcgaaactccgcgggtcggtgagctactcgccgattacttcggcgagaccgaagcccatgcgctgcgcaacgcactggcgcagtcttccgatccaaagcgaccggcgctgatcttcggttggcgcgacggcctgaccggccgcaccttcgacatctcactgcatcgccatgacggtacatcgatcatcgagttcgagcctgcggcggccgaacaggccgacaatccgctgcggctgacgcggcagatcatcgcgcgcaccaaagaactgaagtcgctcgaagagatggccgcacgggtgccgcgctatctgcaggcgatgctcggctatcaccgcgtgatgttgtaccgcttcgcggacgacggctccgggatggtgatcggcgaggcgaagcgcagcgacctcgagagctttctcggtcagcactttccggcgtcgctggtcccgcagcaggcgcggctactgtacttgaagaacgcgatccgcgtggtctcggattcgcgcggcatcagcagccggatcgtgcccgagcacgacgcctccggcgccgcgctcgatctgtcgttcgcgcacctgcgcagcatctcgccctgccatctcgaatttctgcggaacatgggcgtcagcgcctcgatgtcgctgtcgatcatcattgacggcacgctatggggattgatcatctgtcatcattacgagccgcgtgccgtgccgatggcgcagcgcgtcgcggccgaaatgttcgccgacttcttatcgctgcacttcaccgccgcccaccaccaacgc |
| 1802 | CMV | Tom20-LumiFAST | atggtgggtcggaacagcgccatcgccgcgggcgtgtgcggtgccctcttcatagggtactgcatctactttgaccgcaaaagacgaagtgaccccaacttcagatctgccaccATGGAGCATGTTGCCTTTGGCAGTGAGGACATCGAGAACACTCTGGCCAATATGGACGACGAACAACTGGATAGGTTGGCCTTTGGCGTAATTCAGCTCGATGGTGACGGGAATATCCTGCTGTACAATGCTGCTGAAGGGGACATCACTGGCAGAGATCCCAAACAGGTGATTGGGAAGAACTTCTTCAAGGATGTTGCACCTGGAACGGATACTCCCGAGTTTTACGGCAAATTCAAGGAAGGCGCAGCGTCAGGGAATCTGAACACCATGTTCGAATGGACGATACCGACAAGCAGGGGACCAACCAAGGTCAAGGTGCACTTGAAGAAAGCCCTTTCCGGTGACAGATATTGGGTCTTTGTGAAACGGGTGgatattagcggcggcgatattgtcttcacactcgaagatttcgttggggactggcgacagacagccggctacaacctggaccaagtccttgaacagggaggtgtgtccagtttgtttcagaatctcggggtgtccgtaactccgatccaaaggattgtcctgagcggtgaaaatgggctgaagatcgacatccatgtcatcatcccgtatgaaggtctgagcggcgaccaaatgggccagatcgaaaaaatttttaaggtggtgtaccctgtggatgatcatcactttaaggtgatcctgcactatggcacactggtaatcgacggggttacgccgaacatgatcgactatttcggacggccgtatgaaggcatcgccgtgttcgacggcaaaaagatcactgtaacagggaccctgtggaacggcaacaaaattatcgacgagcgcctgatcaaccccgacggctccctgctgttccgagtaaccatcaacggagtgaccggctggcggctgtgcgaacgcattctggcg |
| 1857 | CMV | Lyn11-LumiFAST | ATGggctgcatcaagtccaagggcaaggactccgccggcggcggctccATGGAGCATGTTGCCTTTGGCAGTGAGGACATCGAGAACACTCTGGCCAATATGGACGACGAACAACTGGATAGGTTGGCCTTTGGCGTAATTCAGCTCGATGGTGACGGGAATATCCTGCTGTACAATGCTGCTGAAGGGGACATCACTGGCAGAGATCCCAAACAGGTGATTGGGAAGAACTTCTTCAAGGATGTTGCACCTGGAACGGATACTCCCGAGTTTTACGGCAAATTCAAGGAAGGCGCAGCGTCAGGGAATCTGAACACCATGTTCGAATGGACGATACCGACAAGCAGGGGACCAACCAAGGTCAAGGTGCACTTGAAGAAAGCCCTTTCCGGTGACAGATATTGGGTCTTTGTGAAACGGGTGgatattagcggcggcgatattgtcttcacactcgaagatttcgttggggactggcgacagacagccggctacaacctggaccaagtccttgaacagggaggtgtgtccagtttgtttcagaatctcggggtgtccgtaactccgatccaaaggattgtcctgagcggtgaaaatgggctgaagatcgacatccatgtcatcatcccgtatgaaggtctgagcggcgaccaaatgggccagatcgaaaaaatttttaaggtggtgtaccctgtggatgatcatcactttaaggtgatcctgcactatggcacactggtaatcgacggggttacgccgaacatgatcgactatttcggacggccgtatgaaggcatcgccgtgttcgacggcaaaaagatcactgtaacagggaccctgtggaacggcaacaaaa |

| | | | |
|---|---|---|---|
| | | | ttatcgacgagcgcctgatcaaccccgacggctccctgctgttccgagtaaccatcaacggagtgaccggctggcggctgtgcgaacgcattctggcg |

| Plasmid number | Vector | ORF | |
|---|---|---|---|
| 1601 | pET28 | FRB-DISGGDIS-NanoLuc | atgatcctctggcatgagatgtggcatgaaggcctggaagaggcatctcgtttgtactttggggaaaggaacgtgaaaggcatgtttgaggtgctggagcccttgcatgctatgatggaacggggcccccagactctgaaggaaacatcctttaatcaggcctatggtcgagatttaatggaggcccaagagtggtgcaggaagtacatgaaatcagggaatgtcaaggacctcctccaagcctgggacctctattatcatgtgttccgacgaatctcaaaggatattagcggcggcgatattagcgtcttcacactcgaagatttcgttggggactggcgacagacagccggctacaacctggaccaagtccttgaacagggaggtgtgtccagtttgtttcagaatctcggggtgtccgtaactccgatccaaaggattgtcctgagcggtgaaaatgggctgaagatcgacatccatgtcatcatcccgtatgaaggtctgagcggcgaccaaatgggccagatcgaaaaaatttttaaggtggtgtaccctgtggatgatcatcactttaaggtgatcctgcactatggcacactggtaatcgacggggttacgccgaacatgatcgactatttcggacggccgtatgaaggcatcgccgtgttcgacggcaaaaagatcactgtaacagggaccctgtggaacggcaacaaaattatcgacgagcgcctgatcaaccccgacggctccctgctgttccgagtaaccatcaacggagtgaccggctggcggctgtgcgaacgcattctggcg |
| 1602 | pET28 | FRB-DISGGDIS-pFAST | atgatcctctggcatgagatgtggcatgaaggcctggaagaggcatctcgtttgtactttggggaaaggaacgtgaaaggcatgtttgaggtgctggagcccttgcatgctatgatggaacggggcccccagactctgaaggaaacatcctttaatcaggcctatggtcgagatttaatggaggcccaagagtggtgcaggaagtacatgaaatcagggaatgtcaaggacctcctccaagcctgggacctctattatcatgtgttccgacgaatctcaaaggatattagcggcggcgatattagcATGGAGCATGTTGCCTTTGGCAGTGAGGACATCGAGAACACTCTGGCCAATATGGACGACGAACAACTGGATAGGTTGGCCTTTGGCGTAATTCAGCTCGATGGTGACGGGAATATCCTGCTGTACAATGCTGCTGAAGGGGACATCACTGGCAGAGATCCCAAACAGGTGATTGGGAAGAACTTCTTCAAGGATGTTGCACCTGGAACGGATACTCCCGAGTTTTACGGCAAATTCAAGGAAGGCGCAGCGTCAGGGAATCTGAACACCATGTTCGAATGGACGATACCGACAAGCAGGGGACCAACCAAGGTCAAGGTGCACTTGAAGAAAGCCCTTTCCGGTGACAGATATTGGGTCTTTGTGAAACGGGTG |
| 1603 | pET28 | FKBP-DISGGDIS-NanoLuc | atgGGAGTGCAGGTGGAAACCATCTCCCCAGGAGACGGGCGCACCTTCCCCAAGCGCGGCCAGACCTGCGTGGTGCACTACACCGGGATGCTTGAAGATGGAAAGAAATTTGATTCCTCCCGGGACAGAAACAAGCCCTTTAAGTTTATGCTAGGCAAGCAGGAGGTGATCCGAGGCTGGGAAGAAGGGGTTGCCCAGATGAGTGTGGGTCAGAGAGCCAAACTGACTATATCTCCAGATTATGCCTATGGTGCCACTGGGCACCCAGGCATCATCCCACCACATGCCACTCTCGTCTTCGATGTGGAGCTTCTAAAACTGGAAGAAgatattagcggcggcgatattagcatggtcttcacactcgaagatttcgttggggactggcgacagacagccggctacaacctggaccaagtccttgaacagggaggtgtgtccagtttgtttcagaatctcggggtgtccgtaactccgatccaaaggattgtcctgagcggtgaaaatgggctgaagatcgacatccatgtcatcatcccgtatgaaggtctgagcggcgaccaaatgggccagatcgaaaaaatttttaaggtggtgtaccctgtggatgatcatcactttaaggtgatcctgcactatggcacactggtaatcgacggggttacgccgaacatgatcgactatttcggacggccgtatgaaggcatcgccgtgttcgacggcaaaaagatcactgtaacagggaccctgtggaacggcaacaaaattatcgacgagcgcctgatcaaccccgacggctccctgctgttccgagtaaccatcaacggagtgaccggctggcggctgtgcgaacgcattctggcg |
| 1604 | pET28 | FKBP-DISGGDIS-pFAST | atgGGAGTGCAGGTGGAAACCATCTCCCCAGGAGACGGGCGCACCTTCCCCAAGCGCGGCCAGACCTGCGTGGTGCACTACACCGGGATGCTTGAAGATGGAAAGAAATTTGATTCCTCCCGGGACAGAAACAAGCCCTTTAAGTTTATGCTAGGCAAGCAGGAGGTGATCCGAGGCTGGGAAGAAGGGGTTGCCCAGATGAGTGTGGGTCAGAGAGCCAAACTGACTATATCTCCAGATTATGCCTATGGTGCCACTGGGCACCCAGGCATCATCCCACCACATGCCACTCTCGTCTTCGATGTGGAGCTTCTAAAACTGGAAGAAgatattagcggcggcgatattagcATGGAGCATGTTGCCTTTGGCAGTGAGGACATCGAGAACACTCTGGCCAATATGGACGACGAACAACTGGATAGGTTGGCCTTTGGCGTAATTCAGCTCGATGGTGACGGGAATATCCTGCTGTACAATGCTGCTGAAGGGGACATCACTGGCAGAGATCCCAAACAGGTGATTGGGAAGAACTTCTTCAAGGATGTTGCACCTGGAACGGATACTCCCGAGTTTTACGGCAAATTCAAGGAAGGCGCAGCGTCAGGGAATCTGAACACCATGTTCGAATGGACGATACCGACAAGCAGGGGACCAACCAAGGTCAAGGTGCACTTGAAGAAAGCCCTTTCCGGTGACAGATATTGGGTCTTTGTGAAACGGGTG |
| 1625 | pET28 | pFAST-DENLYFQS-NanoLuc (TEV sensor) | ATGGAGCATGTTGCCTTTGGCAGTGAGGACATCGAGAACACTCTGGCCAATATGGACGACGAACAACTGGATAGGTTGGCCTTTGGCGTAATTCAGCTCGATGGTGACGGGAATATCCTGCTGTACAATGCTGCTGAAGGGGACATCACTGGCAGAGATCCCAAACAGGTGATTGGGAAGAACTTCTTCAAGGATGTTGCACCTGGAACGGATACTCCCGAGTTTTACGGCAAATTCAAGGAAGGCGCAGCGTCAGGGAATCTGAACACCATGTTCGAATGGACGATACCGACAAGCAGGGGACCAACCAAGGTCAAGGTGCACTTGAAGAAAGCCCTTTCCGGTGACAGATATTGGGTCTTTGTGAAACGGGTGgatgaaaacctgtattttcagagcgtcttcacactcgaagatttcgttggggactggcgacagacagccggctacaacctggaccaagtccttgaacagggaggtgtgtccagtttgtttcagaatctcggggtgtccgtaactccgatccaaaggattgtcctgagcggtgaaaatgggctgaagatcgacatccatgtcatcatcccgtatgaaggtctgagcggcgaccaaatgggccagatcgaaaaaatttttaaggtggtgtaccctgtggatgatcatcactttaaggtgatcctgcactatggcacactggtaatcgacggggttacgccgaacatgatcgactatttcggacggccgtatgaaggcatcgccgtgttcgacggcaaaaagatcactgtaacagggaccctgtggaacggcaacaaaattatcgacgagcgcctgatcaaccccgacggctccctgctgttccgagtaaccatcaacggagtgaccggctggcggctgtgcgaacgcattctggcg |

| | | | |
|---|---|---|---|
| 1656 | CMV | pFAST-12-NanoLuc (TEV sensor) | ATGGAGCATGTTGCCTTTGGCAGTGAGGACATCGAGAACACTCTGGCCAA<br>TATGGACGACGAACAACTGGATAGGTTGGCCTTTGGCGTAATTCAGCTCG<br>ATGGTGACGGGAATATCCTGCTGTACAATGCTGCTGAAGGGGACATCACT<br>GGCAGAGATCCCAAACAGGTGATTGGGAAGAACTTCTTCAAGGATGTTGC<br>ACCTGGAACGGATACTCCCGAGTTTTACGGCAAATTCAAGGAAGGCGCAG<br>CGTCAGGGAATCTGAACACCATGTTCGAATGGACGATACCGACAAGCAGG<br>GGACCAACCAAGGTCAAGGTGCACTTGAAGAAAGCCCTTTCCGGTGACAG<br>ATATTGGGTCTTTGTGAAACGGGTGgatattagcgaaaacctgtattttcagagcgatattgtct<br>tcacactcgaagatttcgttggggactggcgacagacagccggctacaacctggaccaagtccttgaacagg<br>gaggtgtgtccagtttgtttcagaatctcggggtgtccgtaactccgatccaaaggattgtcctgagcggtgaaa<br>atgggctgaagatcgacatccatgtcatcatcccgtatgaaggtctgagcggcgaccaaatgggccagatcg<br>aaaaaatttttaaggtggtgtaccctgtggatgatcatcactttaaggtgatcctgcactatggcacactggtaatc<br>gacggggttacgccgaacatgatcgactatttcggacggccgtatgaaggcatcgccgtgttcgacggcaaa<br>aagatcactgtaacagggaccctgtggaacggcaacaaaattatcgacgagcgcctgatcaaccccgacg<br>gctccctgctgttccgagtaaccatcaacggagtgaccggctggcggctgtgcgaacgcattctggcg |
| 1657 | pET28 | pFAST-FRB | atgGAGCATGTTGCCTTTGGCAGTGAGGACATCGAGAACACTCTGGCCAAT<br>ATGGACGACGAACAACTGGATAGGTTGGCCTTTGGCGTAATTCAGCTCGA<br>TGGTGACGGGAATATCCTGCTGTACAATGCTGCTGAAGGGGACATCACTG<br>GCAGAGATCCCAAACAGGTGATTGGGAAGAACTTCTTCAAGGATGTTGCA<br>CCTGGAACGGATACTCCCGAGTTTTACGGCAAATTCAAGGAAGGCGCAGC<br>GTCAGGGAATCTGAACACCATGTTCGAATGGACGATACCGACAAGCAGGG<br>GACCAACCAAGGTCAAGGTGCACTTGAAGAAAGCCCTTTCCGGTGACAGA<br>TATTGGGTCTTTGTGAAACGGGTGatcctctggcatgagatgtggcatgaaggcctggaaga<br>ggcatctcgtttgtactttggggaaaggaacgtgaaaggcatgtttgaggtgctggagcccttgcatgctatgat<br>ggaacggggcccccagactctgaaggaaacatcctttaatcaggcctatggtcgagatttaatggaggccca<br>agagtggtgcaggaagtacatgaaatcagggaatgtcaaggacctcctccaagcctgggacctctattatcat<br>gtgttccgacgaatctcaaag |
| 1658 | pET28 | pFAST-DI-FRB | atgGAGCATGTTGCCTTTGGCAGTGAGGACATCGAGAACACTCTGGCCAAT<br>ATGGACGACGAACAACTGGATAGGTTGGCCTTTGGCGTAATTCAGCTCGA<br>TGGTGACGGGAATATCCTGCTGTACAATGCTGCTGAAGGGGACATCACTG<br>GCAGAGATCCCAAACAGGTGATTGGGAAGAACTTCTTCAAGGATGTTGCA<br>CCTGGAACGGATACTCCCGAGTTTTACGGCAAATTCAAGGAAGGCGCAGC<br>GTCAGGGAATCTGAACACCATGTTCGAATGGACGATACCGACAAGCAGGG<br>GACCAACCAAGGTCAAGGTGCACTTGAAGAAAGCCCTTTCCGGTGACAGA<br>TATTGGGTCTTTGTGAAACGGGTGgatattatcctctggcatgagatgtggcatgaaggcctg<br>gaagaggcatctcgtttgtactttggggaaaggaacgtgaaaggcatgtttgaggtgctggagcccttgcatgc<br>tatgatggaacggggcccccagactctgaaggaaacatcctttaatcaggcctatggtcgagatttaatggag<br>gcccaagagtggtgcaggaagtacatgaaatcagggaatgtcaaggacctcctccaagcctgggacctcta<br>ttatcatgtgttccgacgaatctcaaag |
| 1659 | pET28 | FKBP-NanoLuc | atgGGAGTGCAGGTGGAAACCATCTCCCCAGGAGACGGGCGCACCTTCCC<br>CAAGCGCGGCCAGACCTGCGTGGTGCACTACACCGGGATGCTTGAAGAT<br>GGAAAGAAATTTGATTCCTCCCGGGACAGAAACAAGCCCTTTAAGTTTATG<br>CTAGGCAAGCAGGAGGTGATCCGAGGCTGGGAAGAAGGGGTTGCCCAGA<br>TGAGTGTGGGTCAGAGAGCCAAACTGACTATATCTCCAGATTATGCCTATG<br>GTGCCACTGGGCACCCAGGCATCATCCCACCACATGCCACTCTCGTCTTC<br>GATGTGGAGCTTCTAAAACTGGAAGAAatggtcttcacactcgaagatttcgttggggactgg<br>cgacagacagccggctacaacctggaccaagtccttgaacagggaggtgtgtccagtttgtttcagaatctcg<br>gggtgtccgtaactccgatccaaaggattgtcctgagcggtgaaaatgggctgaagatcgacatccatgtcat<br>catcccgtatgaaggtctgagcggcgaccaaatgggccagatcgaaaaaatttttaaggtggtgtaccctgtg<br>gatgatcatcactttaaggtgatcctgcactatggcacactggtaatcgacggggttacgccgaacatgatcga<br>ctatttcggacggccgtatgaaggcatcgccgtgttcgacggcaaaaagatcactgtaacagggaccctgtg<br>gaacggcaacaaaattatcgacgagcgcctgatcaaccccgacggctccctgctgttccgagtaaccatcaa<br>cggagtgaccggctggcggctgtgcgaacgcattctggcg |
| 1660 | pET28 | FKBP-DI-NanoLuc | atgGGAGTGCAGGTGGAAACCATCTCCCCAGGAGACGGGCGCACCTTCCC<br>CAAGCGCGGCCAGACCTGCGTGGTGCACTACACCGGGATGCTTGAAGAT<br>GGAAAGAAATTTGATTCCTCCCGGGACAGAAACAAGCCCTTTAAGTTTATG<br>CTAGGCAAGCAGGAGGTGATCCGAGGCTGGGAAGAAGGGGTTGCCCAGA<br>TGAGTGTGGGTCAGAGAGCCAAACTGACTATATCTCCAGATTATGCCTATG<br>GTGCCACTGGGCACCCAGGCATCATCCCACCACATGCCACTCTCGTCTTC<br>GATGTGGAGCTTCTAAAACTGGAAGAAgatattatggtcttcacactcgaagatttcgttgggg<br>actggcgacagacagccggctacaacctggaccaagtccttgaacagggaggtgtgtccagtttgtttcaga<br>atctcggggtgtccgtaactccgatccaaaggattgtcctgagcggtgaaaatgggctgaagatcgacatcca<br>tgtcatcatcccgtatgaaggtctgagcggcgaccaaatgggccagatcgaaaaaatttttaaggtggtgtacc<br>ctgtggatgatcatcactttaaggtgatcctgcactatggcacactggtaatcgacggggttacgccgaacatg<br>atcgactatttcggacggccgtatgaaggcatcgccgtgttcgacggcaaaaagatcactgtaacagggacc<br>ctgtggaacggcaacaaaattatcgacgagcgcctgatcaaccccgacggctccctgctgttccgagtaacc<br>atcaacggagtgaccggctggcggctgtgcgaacgcattctggcg |
| 1670 | pET28 | pFAST-19-NanoLuc (TEV sensor) | ATGGAGCATGTTGCCTTTGGCAGTGAGGACATCGAGAACACTCTGGCCAA<br>TATGGACGACGAACAACTGGATAGGTTGGCCTTTGGCGTAATTCAGCTCG<br>ATGGTGACGGGAATATCCTGCTGTACAATGCTGCTGAAGGGGACATCACT<br>GGCAGAGATCCCAAACAGGTGATTGGGAAGAACTTCTTCAAGGATGTTGC<br>ACCTGGAACGGATACTCCCGAGTTTTACGGCAAATTCAAGGAAGGCGCAG<br>CGTCAGGGAATCTGAACACCATGTTCGAATGGACGATACCGACAAGCAGG<br>GGACCAACCAAGGTCAAGGTGCACTTGAAGAAAGCCCTTTCCGGTGACAG<br>ATATTGGGTCTTTGTGAAACGGGTGgatattagcggcggcgatgaaaacctgtattttcagag |

| | | | |
|---|---|---|---|
| | | | cgatattagcggcggcgatgtcttcacactcgaagatttcgttggggactggcgacagacagccggctacaacctggaccaagtccttgaacagggaggtgtgtccagtttgtttcagaatctcggggtgtccgtaactccgatccaaaggattgtcctgagcggtgaaaatgggctgaagatcgacatccatgtcatcatcccgtatgaaggtctgagcggcgaccaaatgggccagatcgaaaaaatttttaaggtggtgtaccctgtggatgatcatcactttaaggtgatcctgcactatggcacactggtaatcgacggggttacgccgaacatgatcgactatttcggacggccgtatgaaggcatcgccgtgttcgacggcaaaaagatcactgtaacagggaccctgtggaacggcaacaaaattatcgacgagcgcctgatcaaccccgacggctccctgctgttccgagtaaccatcaacggagtgaccggctggcggctgtgcgaacgcattctggcg |
| 1703 | pET28 | NanoLuc-DISGGDIS-FRB | atggtcttcacactcgaagatttcgttggggactggcgacagacagccggctacaacctggaccaagtccttgaacagggaggtgtgtccagtttgtttcagaatctcggggtgtccgtaactccgatccaaaggattgtcctgagcggtgaaaatgggctgaagatcgacatccatgtcatcatcccgtatgaaggtctgagcggcgaccaaatgggccagatcgaaaaaatttttaaggtggtgtaccctgtggatgatcatcactttaaggtgatcctgcactatggcacactggtaatcgacggggttacgccgaacatgatcgactatttcggacggccgtatgaaggcatcgccgtgttcgacggcaaaaagatcactgtaacagggaccctgtggaacggcaacaaaattatcgacgagcgcctgatcaaccccgacggctccctgctgttccgagtaaccatcaacggagtgaccggctggcggctgtgcgaacgcattctggcggatattagcggcggcgatattagcatcctctggcatgagatgtggcatgaaggcctggaagaggcatctcgtttgtactttggggaaaggaacgtgaaaggcatgtttgaggtgctggagcccttgcatgctatgatggaacggggcccccagactctgaaggaaacatcctttaatcaggcctatggtcgagatttaatggaggcccaagagtggtgcaggaagtacatgaaatcagggaatgtcaaggacctcctccaagcctgggacctctattatcatgtgttccgacgaatctcaaag |
| 1704 | pET28 | pFAST-DISGGDIS-FRB | atgGAGCATGTTGCCTTTGGCAGTGAGGACATCGAGAACACTCTGGCCAATATGGACGACGAACAACTGGATAGGTTGGCCTTTGGCGTAATTCAGCTCGATGGTGACGGGAATATCCTGCTGTACAATGCTGCTGAAGGGGACATCACTGGCAGAGATCCCAAACAGGTGATTGGGAAGAACTTCTTCAAGGATGTTGCACCTGGAACGGATACTCCCGAGTTTTACGGCAAATTCAAGGAAGGCGCAGCGTCAGGGAATCTGAACACCATGTTCGAATGGACGATACCGACAAGCAGGGGACCAACCAAGGTCAAGGTGCACTTGAAGAAAGCCCTTTCCGGTGACAGATATTGGGTCTTTGTGAAACGGGTGgatattagcggcggcgatattagcatcctctggcatgagatgtggcatgaaggcctggaagaggcatctcgtttgtactttggggaaaggaacgtgaaaggcatgtttgaggtgctggagcccttgcatgctatgatggaacggggcccccagactctgaaggaaacatcctttaatcaggcctatggtcgagatttaatggaggcccaagagtggtgcaggaagtacatgaaatcagggaatgtcaaggacctcctccaagcctgggacctctattatcatgtgttccgacgaatctcaaag |
| 1705 | pET28 | NanoLuc-DISGGDIS-FKBP | atgatggtcttcacactcgaagatttcgttggggactggcgacagacagccggctacaacctggaccaagtccttgaacagggaggtgtgtccagtttgtttcagaatctcggggtgtccgtaactccgatccaaaggattgtcctgagcggtgaaaatgggctgaagatcgacatccatgtcatcatcccgtatgaaggtctgagcggcgaccaaatgggccagatcgaaaaaatttttaaggtggtgtaccctgtggatgatcatcactttaaggtgatcctgcactatggcacactggtaatcgacggggttacgccgaacatgatcgactatttcggacggccgtatgaaggcatcgccgtgttcgacggcaaaaagatcactgtaacagggaccctgtggaacggcaacaaaattatcgacgagcgcctgatcaaccccgacggctccctgctgttccgagtaaccatcaacggagtgaccggctggcggctgtgcgaacgcattctggcggatattagcggcggcgatattagcGGAGTGCAGGTGGAAACCATCTCCCCAGGAGACGGGCGCACCTTCCCCAAGCGCGGCCAGACCTGCGTGGTGCACTACACCGGGATGCTTGAAGATGGAAAGAAATTTGATTCCTCCCGGGACAGAAACAAGCCCTTTAAGTTTATGCTAGGCAAGCAGGAGGTGATCCGAGGCTGGGAAGAAGGGGTTGCCCAGATGAGTGTGGGTCAGAGAGCCAAACTGACTATATCTCCAGATTATGCCTATGGTGCCACTGGGCACCCAGGCATCATCCCACCACATGCCACTCTCGTCTTCGATGTGGAGCTTCTAAAACTGGAAGAA |
| 1706 | pET28 | pFAST-DISGGDIS-FKBP | atgGAGCATGTTGCCTTTGGCAGTGAGGACATCGAGAACACTCTGGCCAATATGGACGACGAACAACTGGATAGGTTGGCCTTTGGCGTAATTCAGCTCGATGGTGACGGGAATATCCTGCTGTACAATGCTGCTGAAGGGGACATCACTGGCAGAGATCCCAAACAGGTGATTGGGAAGAACTTCTTCAAGGATGTTGCACCTGGAACGGATACTCCCGAGTTTTACGGCAAATTCAAGGAAGGCGCAGCGTCAGGGAATCTGAACACCATGTTCGAATGGACGATACCGACAAGCAGGGGACCAACCAAGGTCAAGGTGCACTTGAAGAAAGCCCTTTCCGGTGACAGATATTGGGTCTTTGTGAAACGGGTGgatattagcggcggcgatattagcGGAGTGCAGGTGGAAACCATCTCCCCAGGAGACGGGCGCACCTTCCCCAAGCGCGGCCAGACCTGCGTGGTGCACTACACCGGGATGCTTGAAGATGGAAAGAAATTTGATTCCTCCCGGGACAGAAACAAGCCCTTTAAGTTTATGCTAGGCAAGCAGGAGGTGATCCGAGGCTGGGAAGAAGGGGTTGCCCAGATGAGTGTGGGTCAGAGAGCCAAACTGACTATATCTCCAGATTATGCCTATGGTGCCACTGGGCACCCAGGCATCATCCCACCACATGCCACTCTCGTCTTCGATGTGGAGCTTCTAAAACTGGAAGAA |
| 1797 | pET28 | pFAST-12-NanoLuc (TEV sensor) | ATGGAGCATGTTGCCTTTGGCAGTGAGGACATCGAGAACACTCTGGCCAATATGGACGACGAACAACTGGATAGGTTGGCCTTTGGCGTAATTCAGCTCGATGGTGACGGGAATATCCTGCTGTACAATGCTGCTGAAGGGGACATCACTGGCAGAGATCCCAAACAGGTGATTGGGAAGAACTTCTTCAAGGATGTTGCACCTGGAACGGATACTCCCGAGTTTTACGGCAAATTCAAGGAAGGCGCAGCGTCAGGGAATCTGAACACCATGTTCGAATGGACGATACCGACAAGCAGGGGACCAACCAAGGTCAAGGTGCACTTGAAGAAAGCCCTTTCCGGTGACAGATATTGGGTCTTTGTGAAACGGGTGgatattagcgaaaacctgtattttcagagcgatattgtcttcacactcgaagatttcgttggggactggcgacagacagccggctacaacctggaccaagtccttgaacagggaggtgtgtccagtttgtttcagaatctcggggtgtccgtaactccgatccaaaggattgtcctgagcggtgaaaatgggctgaagatcgacatccatgtcatcatcccgtatgaaggtctgagcggcgaccaaatgggccagatcgaaaaaatttttaaggtggtgtaccctgtggatgatcatcactttaaggtgatcctgcactatggcacactggtaatcgacggggttacgccgaacatgatcgactatttcggacggccgtatgaaggcatcgccgtgttcgacggcaaa |

| | | | |
|---|---|---|---|
| | | | aagatcactgtaacagggaccctgtggaacggcaacaaaattatcgacgagcgcctgatcaaccccgacggctccctgctgttccgagtaaccatcaacggagtgaccggctggcggctgtgcgaacgcattctggcg |
| 1821 | CMV | pFAST-FRB | atgGAGCATGTTGCCTTTGGCAGTGAGGACATCGAGAACACTCTGGCCAATATGGACGACGAACAACTGGATAGGTTGGCCTTTGGCGTAATTCAGCTCGATGGTGACGGGAATATCCTGCTGTACAATGCTGCTGAAGGGGACATCACTGGCAGAGATCCCAAACAGGTGATTGGGAAGAACTTCTTCAAGGATGTTGCACCTGGAACGGATACTCCCGAGTTTTACGGCAAATTCAAGGAAGGCGCAGCGTCAGGGAATCTGAACACCATGTTCGAATGGACGATACCGACAAGCAGGGGACCAACCAAGGTCAAGGTGCACTTGAAGAAAGCCCTTTCCGGTGACAGATATTGGGTCTTTGTGAAACGGGTGatcctctggcatgagatgtggcatgaaggcctggaagaggcatctcgtttgtactttggggaaaggaacgtgaaaggcatgtttgaggtgctggagcccttgcatgctatgatggaacggggcccccagactctgaaggaaacatcctttaatcaggcctatggtcgagatttaatggaggcccaagagtggtgcaggaagtacatgaaatcagggaatgtcaaggacctcctccaagcctgggacctctattatcatgtgttccgacgaatctcaaag |
| 1822 | CMV | FKBP-NanoLuc | atgGGAGTGCAGGTGGAAACCATCTCCCCAGGAGACGGGCGCACCTTCCCCAAGCGCGGCCAGACCTGCGTGGTGCACTACACCGGGATGCTTGAAGATGGAAAGAAATTTGATTCCTCCCGGGACAGAAACAAGCCCTTTAAGTTTATGCTAGGCAAGCAGGAGGTGATCCGAGGCTGGGAAGAAGGGGTTGCCCAGATGAGTGTGGGTCAGAGAGCCAAACTGACTATATCTCCAGATTATGCCTATGGTGCCACTGGGCACCCAGGCATCATCCCACCACATGCCACTCTCGTCTTCGATGTGGAGCTTCTAAAACTGGAAGAAatggtcttcacactcgaagatttcgttggggactggcgacagacagccggctacaacctggaccaagtccttgaacagggaggtgtgtccagtttgtttcagaatctcggggtgtccgtaactccgatccaaaggattgtcctgagcggtgaaaatgggctgaagatcgacatccatgtcatcatcccgtatgaaggtctgagcggcgaccaaatgggccagatcgaaaaaatttttaaggtggtgtaccctgtggatgatcatcactttaaggtgatcctgcactatggcacactggtaatcgacggggttacgccgaacatgatcgactatttcggacggccgtatgaaggcatcgccgtgttcgacggcaaaaagatcactgtaacagggaccctgtggaacggcaacaaaattatcgacgagcgcctgatcaaccccgacggctccctgctgttccgagtaaccatcaacggagtgaccggctggcggctgtgcgaacgcattctggcg |

### Supplementary References


(1) Benaissa, H.; Ounoughi, K.; Aujard, I.; Fischer, E.; Goïame, R.; Nguyen, J.; Tebo, A. G.; Li, C.; Le Saux, T.; Bertolin, G.; Tramier, M.; Danglot, L.; Pietrancosta, N.; Morin, X.; Jullien, L.; Gautier, A. Engineering of a Fluorescent Chemogenetic Reporter with Tunable Color for Advanced Live-Cell Imaging. *Nat. Commun.* **2021**, *12* (1), 6989.

(2) Plamont, M.-A.; Billon-Denis, E.; Maurin, S.; Gauron, C.; Pimenta, F. M.; Specht, C. G.; Shi, J.; Quérard, J.; Pan, B.; Rossignol, J.; Moncoq, K.; Morellet, N.; Volovitch, M.; Lescop, E.; Chen, Y.; Triller, A.; Vriz, S.; Le Saux, T.; Jullien, L.; Gautier, A. Small Fluorescence-Activating and Absorption-Shifting Tag for Tunable Protein Imaging in Vivo. *Proc. Natl. Acad. Sci. U. S. A.* **2016**, *113* (3), 497–502.

(3) Li, C.; Plamont, M.-A.; Sladitschek, H. L.; Rodrigues, V.; Aujard, I.; Neveu, P.; Le Saux, T.; Jullien, L.; Gautier, A. Dynamic Multicolor Protein Labeling in Living Cells. *Chem. Sci.* **2017**, *8* (8), 5598–5605.

(4) Mineev, K. S.; Goncharuk, S. A.; Goncharuk, M. V.; Povarova, N. V.; Sokolov, A. I.; Baleeva, N. S.; Smirnov, A. Y.; Myasnyanko, I. N.; Ruchkin, D. A.; Bukhdruker, S.; Remeeva, A.; Mishin, A.; Borshchevskiy, V.; Gordeliy, V.; Arseniev, A. S.; Gorbachev, D. A.; Gavrikov, A. S.; Mishin, A. S.; Baranov, M. S. NanoFAST: Structure-Based Design of a Small Fluorogen-Activating Protein with Only 98 Amino Acids. *Chem. Sci.* **2021**, *12*, 6719–6725.

(5) Gibson, D. G.; Young, L.; Chuang, R. Y.; Venter, J. C.; Hutchison, C. A.; Smith, H. O. Enzymatic Assembly of DNA Molecules up to Several Hundred Kilobases. *Nat. Methods* **2009**, *6*, 343–345.

(6) Broch, F.; El Hajji, L.; Pietrancosta, N.; Gautier, A. Engineering of Tunable Allosteric-like Fluorogenic Protein Sensors. *ACS Sens.* **2023**, *8* (10), 3933–3942.

(7) Hiblot, J.; Yu, Q.; Sabbadini, M. D. B.; Reymond, L.; Xue, L.; Schena, A.; Sallin, O.; Hill, N.; Griss, R.; Johnsson, K. Luciferases with Tunable Emission Wavelengths. *Angew. Chem. Int. Ed Engl.* **2017**, *56* (46), 14556–14560.

(8) El Hajji, L.; Bunel, B.; Joliot, O.; Li, C.; Tebo, A. G.; Rampon, C.; Volovitch, M.; Fischer, E.; Pietrancosta, N.; Perez, F.; Morin, X.; Vriz, S.; Gautier, A. A Tunable and Versatile Chemogenetic Near-Infrared Fluorescent Reporter. *Nat. Commun.* **2025**, *16* (1), 2594.
